# A single bacterial genus maintains root development in a complex microbiome

**DOI:** 10.1101/645655

**Authors:** Omri M. Finkel, Isai Salas-González, Gabriel Castrillo, Jonathan M. Conway, Theresa F. Law, Paulo José Pereira Lima Teixeira, Ellie D. Wilson, Connor R. Fitzpatrick, Corbin D. Jones, Jeffery L. Dangl

**Author notes:** These authors contributed equally to this work (Omri M. Finkel, Isai Salas-González, Gabriel Castrillo, Jonathan M. Conway). Future Food Beacon of Excellence and the School of Biosciences, University of Nottingham, Sutton Bonington, United Kingdom (Gabriel Castrillo). Department of Biology, “Luiz de Queiroz” College of Agriculture (ESALQ), University of São Paulo (USP), Piracicaba, São Paulo, Brazil.

## Abstract

Plants grow within a complex web of species interacting with each other and with the plant. Many of these interactions are governed by a wide repertoire of chemical signals, and the resulting chemical landscape of the rhizosphere can strongly affect root health and development. To understand how microbe-microbe interactions influence root development in Arabidopsis, we established a model system for plant-microbe-microbe-environment interactions. We inoculated seedlings with a 185-member bacterial synthetic community (SynCom), manipulated the abiotic environment, and measured bacterial colonization of the plant. This enabled classification of the SynCom into four modules of co-occurring strains. We deconstructed the SynCom based on these modules, identifying microbe-microbe interactions that determine root phenotypes. These interactions primarily involve a single bacterial genus, *Variovorax*, which completely reverts severe root growth inhibition (RGI) induced by a wide diversity of bacterial strains as well as by the entire 185-member community. We demonstrate that *Variovorax* manipulate plant hormone levels to balance this ecologically realistic root community’s effects on root development. We identify a novel auxin degradation operon in the *Variovorax* genome that is necessary and sufficient for RGI reversion. Therefore, metabolic signal interference shapes bacteria-plant communication networks and is essential for maintaining the root’s developmental program. Optimizing the feedbacks that shape chemical interaction networks in the rhizosphere provides a promising new ecological strategy towards the development of more resilient and productive crops.

## Main

Plant phenotypes, and ultimately fitness, are influenced by the microbes living in close association with them^1–4^. These microbes, collectively termed the plant microbiota, assemble based on plant- and environmentally-derived cues^5, 6^ resulting in myriad plant-microbe interactions. Beneficial and detrimental microbial effects on plants can be either direct^1, 7–10^, or an indirect consequence of microbe-microbe interactions^3, 11^. The contribution of microbe-microbe interactions within the microbiota to community assembly and to the community’s effect on the host is unknown, and it is therefore unclear to what extent binary plant-microbe interactions hold in complex ecological contexts. While antagonistic microbe-microbe interactions are known to play an important role in shaping plant microbiota and protecting plants from pathogens^3^, another potentially significant class of interactions is metabolic signal interference^8, 12^: rather than direct antagonism, microbes interfere with the delivery of chemical signals produced by other microbes, altering plant-microbe signaling^13–15^.

Plant hormones, in particular auxins, are both produced and degraded by an abundance of plant-associated microbes^16–19^. Microbially-derived auxins can have effects on plants ranging from growth-promoting to disease-inducing, depending on context and concentration^20^. The plant’s intrinsic root developmental patterns are dependent on finely calibrated auxin and ethylene concentration gradients with fine differences across tissues and cell types^21^, and it is not known how the plant integrates exogenous, microbially-derived auxin fluxes into its developmental plan.

Here we apply a synthetic community (SynCom) that is reasonably representative of wild soil root-associated microbiomes, to axenic plants to ask how microbe-microbe interactions shape plant root development. We use plant colonization patterns across 16 abiotic conditions to guide stepwise deconstruction of the SynCom, leading to the identification of multiple levels of microbe-microbe interactions that interfere with the additivity of bacterial effects on root development. We demonstrate that a single bacterial genus, *Variovorax*, is required for maintaining the root’s intrinsically controlled developmental program by tuning its chemical landscape. Finally, we identify the locus conserved across *Variovorax* strains that is responsible for this phenotype.

### Consistent community assembly across environmental conditions

To model plant-microbiota interactions in a fully controlled setting, we established a plant-microbiota microcosm representing the native bacterial root-derived microbiota on agar plates. We inoculated 7-day-old seedlings with a defined 185-member bacterial SynCom (Extended Data Fig. 1a, Supplementary Table 1) composed of genome-sequenced isolates obtained from surface sterilized Arabidopsis roots (Methods 1). The composition of this SynCom captures the diversity of Actinobacteria, Proteobacteria and Bacteroidetes; the three major root-enriched phyla^6, 22, 23^; and Firmicutes, which are abundant in plant-associated culture collections^24^. To test the robustness of microbiota assembly to the abiotic environment, we exposed this microcosm to each of 16 different abiotic contexts by manipulating one of four variables (salinity, temperature, phosphate concentration, and pH). We measured SynCom composition in root, shoot and agar fractions 12 days post-inoculation using 16S rRNA amplicon sequencing.

The composition of the resulting root and shoot microbiota recapitulated phylum-level plant enrichment patterns seen in soil-planted *Arabidopsis thaliana*, with comparable proportions of Proteobacteria, Bacteroidetes and Actinobacteria (Extended Data Fig. 1b). Firmicutes, which are not plant-enriched, were reduced to <0.1% of the relative abundance. We observed similar patterns in seedlings grown in sterilized potting soil^25^ inoculated with the same SynCom (Methods 2). Both relative abundances and plant enrichment patterns at the unique sequence (USeq)-levels were significantly correlated between the agar- and soil-based systems, confirming the applicability of the relatively high-throughput agar-based system as a model for plant microbiota assembly (Extended Data Fig. 1c).

Fraction (substrate, root, shoot) (Extended Data Fig. 1d), explained most (40%) of the variance across all abiotic variables. Abiotic conditions significantly affected both alpha- (Extended Data Fig. 1e) and beta-diversity (Fig. 1a). We calculated pairwise correlations in relative abundance across all samples, and identified four well-defined modules of co-occurring strains: A, B, C and D (Fig. 1b, Supplementary Table 2). These modules formed distinct phylogenetically-structured guilds in association with the plant: module A contained mainly Gammaproteobacteria and was predominantly more abundant in the substrate than in the seedling; module B contained mainly low-abundance Firmicutes, with no significant seedling enrichment trend; modules C and D were composed mainly of Alphaproteobacteria and Actinobacteria, respectively, and showed plant-enrichment across all abiotic conditions (Figure 1a, Supplementary Table 2). Both Alphaproteobacteria (module C) and Actinobacteria (module D) are consistently plant-enriched across plant species^23^, suggesting that these clades contain plant-association traits that are deeply rooted in their evolutionary histories.

**Fig. 1.**
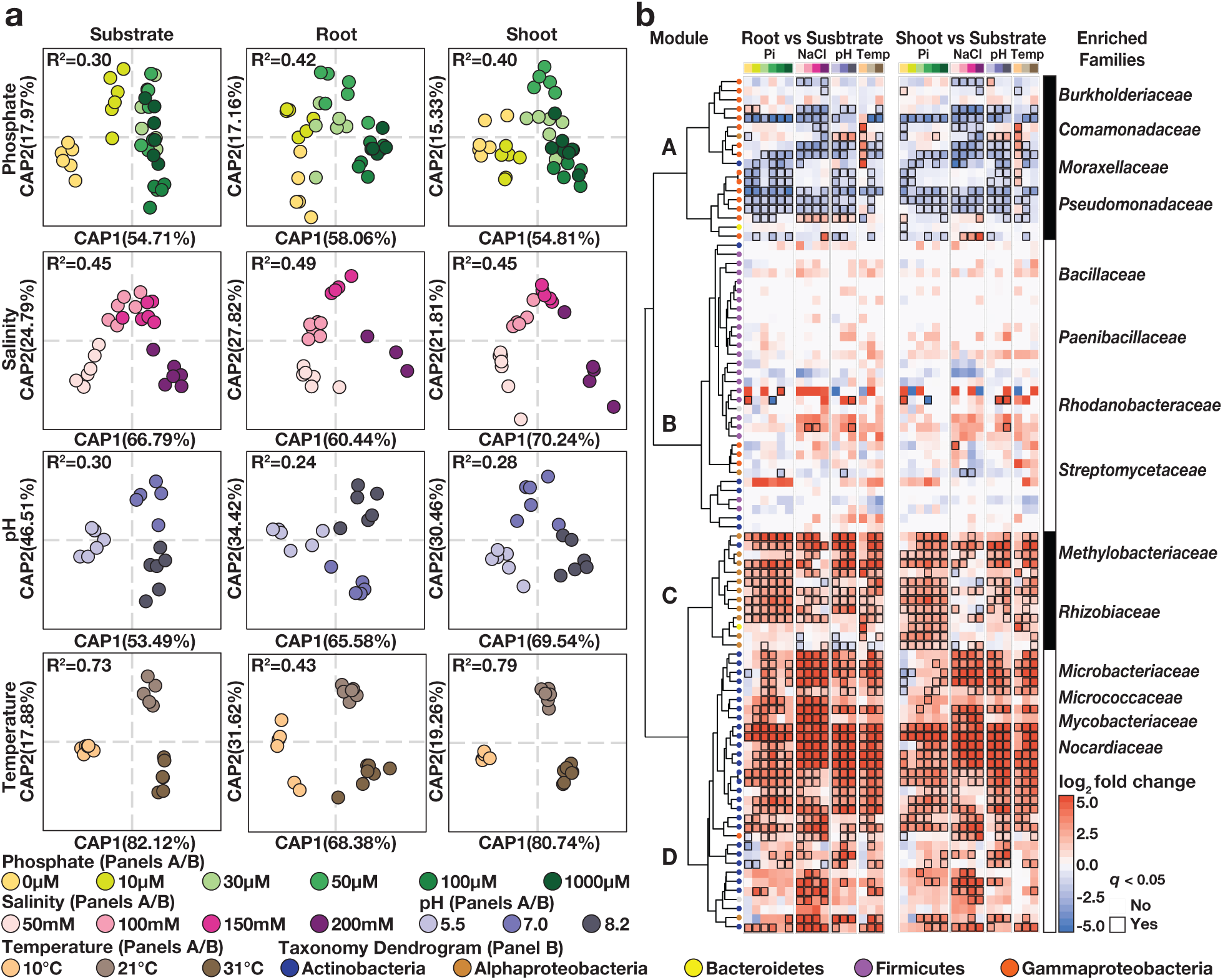
Reproducible effects of abiotic conditions on the synthetic community assembly. **(a)** Canonical analysis of principal coordinates (CAP) scatterplots showing the influence of each of the four abiotic gradients (phosphate, salinity, pH, temperature) within substrate, root and shoot fractions. PERMANOVA R2 values are shown within each plot. **(b)** Fraction enrichment patterns of the SynCom across abiotic gradients. Each row represents a USeq. Letters on the dendrogram represent the four modules of co-occurring strains (A, B, C, D). Dendrogram tips are colored by taxonomy. The heatmaps are colored by log2 fold changes derived from a fitted GLM. Positive fold changes (red gradient) represent enrichments in plant tissue (root or shoot) compared with substrate, negative fold changes (blue gradient) represent depletion in plant tissue compared with substrate. Comparisons with q-value < 0.05 are contoured in black. Family bar highlighenriched families within each module.

### Root development is controlled by multiple microbe-microbe interactions

We next asked whether the different modules of co-occurring strains play different roles in determining plant phenotypes (Methods 3). We inoculated seedlings with SynComs composed of modules A, B, C and D singly, or in all six possible pairwise module combinations, and imaged the seedlings 12 days post-inoculation. We observed strong primary root growth arrest in seedlings inoculated with plant-enriched modules C or D (Fig. 2a, c). Root growth inhibition (RGI) did not occur in seedlings inoculated with modules A or B, which do not contain plant-enriched strains (Fig. 2a, Supplementary Table 3). To test whether the root phenotype derived from each module is an additive outcome of its individual constituents, we inoculated seedlings in mono-association with each of the 185 SynCom members (Methods 4). We observed that 34 taxonomically diverse strains, distributed across all four modules, induced RGI (Fig. 2b, Extended Data Fig. 2, Supplementary Table 4). However, neither the full SynCom nor derived SynComs consisting of modules A or B exhibited RGI (Fig. 2a). Thus, binary plant-microbe interactions were not predictive of interactions in this complex community context.

**Fig. 2.**
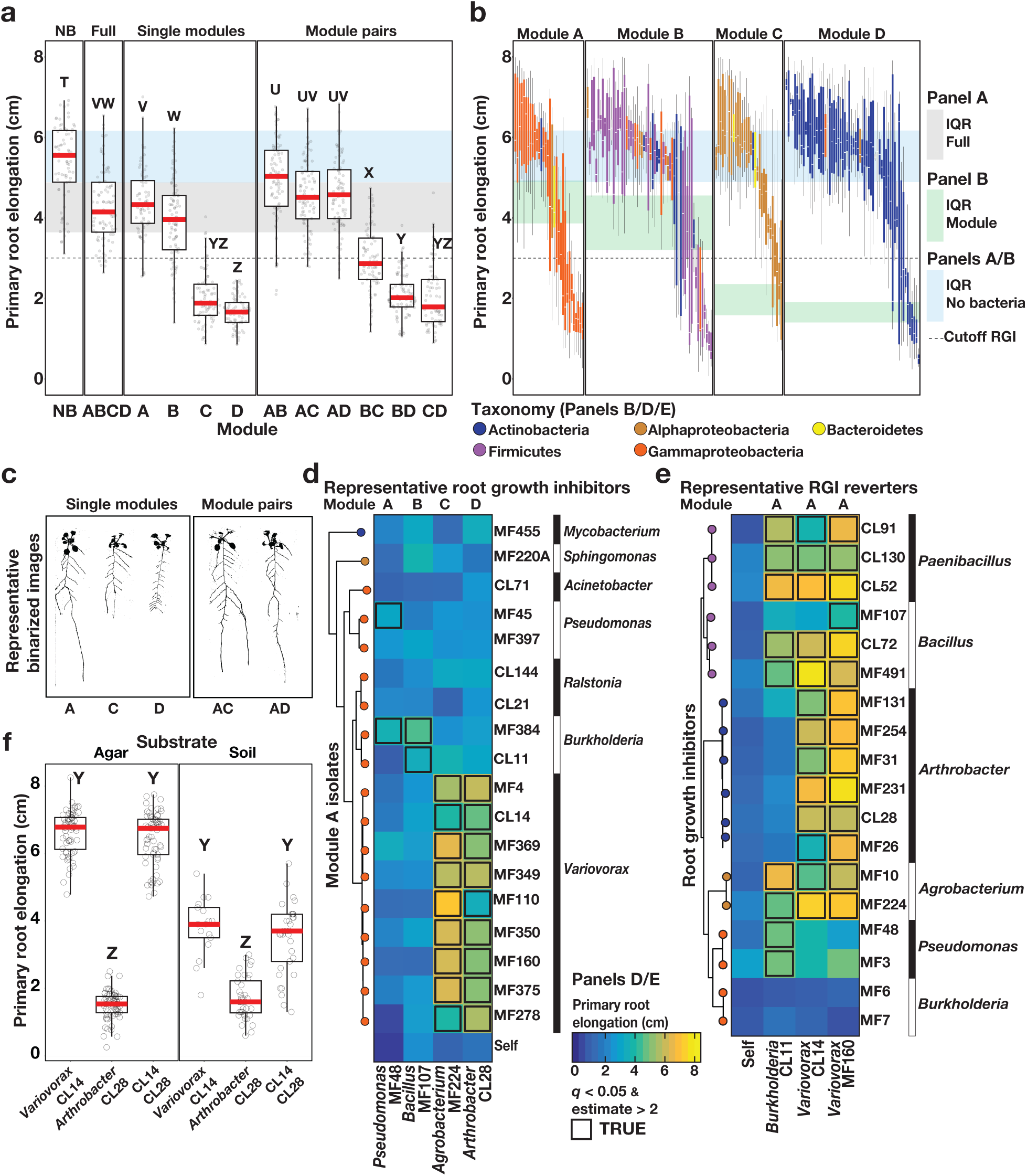
Arabidopsis root length is governed by multiple bacteria-bacteria interactions within a community. **(a)** Primary root elongation of seedlings grown with no bacteria (NB), with the full 185-member SynCom (Full) or with its subsets: Modules A, B, C and D alone (single modules), as well as all six possible pairwise combination of modules (module pairs). Differences between treatments are denoted using the compact letter display. **(b)** Primary root elongation of seedlings inoculated with single bacterial isolates. Isolates are colored by taxonomy and grouped by module membership. The strips across the panels correspond to the interquartile range (IQR) as noted at far right. The dotted line represents the cutoff used to classify isolates as root growth inhibiting (cutoff RGI). **(c)** Binarized image of representative seedlings inoculated with modules A, C and D, and with module combinations AC and AD. (d, e) Heatmaps colored by average primary root elongation of seedlings inoculated with different pairs of bacterial isolates: **(d)** with four representative RGI-inducing strains from each module (columns) alone (Self) or in combination with isolates from module A (rows); **(e)** with eighteen RGI-inducing strains (rows) alone (Self) or in combination with Burkholderia CL11, Variovorax CL14 or Variovorax MF160 (columns). Statistically significant RGI reversions are contoured in black. **(f)** Primary root elongation of uninoculated seedlings (NB) or seedlings inoculated with Arthrobacter CL28 and Variovorax CL14 isolates individually or jointly. Letters indicate post-hoc significance.

In seedlings inoculated with module pairs, we observed an epistatic interaction: in the presence of module A, RGI caused by modules C and D was reverted (Fig. 2a). Thus, through deconstructing the SynCom into four modules, we found that bacterial effects on root development are governed by multiple levels of microbe-microbe interactions. This is exemplified by at least four instances: within modules A or B and between module A and modules C or D. Since three of these interactions involve module A, we predicted that this module contains strains that strongly attenuate RGI, preserving stereotypic root development.

### *Variovorax* are necessary and sufficient to maintain stereotypic plant root development within complex microbiota

To identify strains within module A responsible for intra- and inter-module RGI attenuation, we reduced our system to a tripartite plant-microbe-microbe system (Methods 5). We individually screened the 18 non-RGI strains from module A for their ability to attenuate RGI caused by representative strains from all four modules. We found that all strains from a single genus, *Variovorax* (Family Comamonadaceae), suppressed RGI caused by representative RGI-inducing strains from module C (*Agrobacterium* MF224) and module D (*Arthrobacter* CL28; Fig. 2d, Supplementary Table 5). The strains from modules A (*Pseudomonas* MF48) and B (*Bacillus* MF107), were not suppressed by *Variovorax*, but rather by two closely related *Burkholderia* strains (CL11, MF384). A similar pattern was observed when we screened three selected RGI-suppressing *Variovorax* strains (CL14, MF160) and *Burkholderia* CL11, against a diverse set of RGI-inducers. *Variovorax* attenuated 13 of the 18 RGI-inducers tested (Fig. 2e, Supplementary Table 5).

To test whether RGI induction and repression observed on agar occur in soil as well, we germinated Arabidopsis on sterile soil inoculated with an RGI-suppressor-inducer pair composed of the RGI-inducer *Arthrobacter* CL28 and the RGI-attenuator *Variovorax* CL14 (Methods 2). As expected, *Arthrobacter* CL28 induced RGI which was reverted by *Variovorax* CL14 in soil (Fig. 2f, Supplementary Table 6). We generalized this observation by showing that *Variovorax*-mediated RGI attenuation extended to tomato seedlings, where *Variovorax* CL14 reverted *Arthrobacter* CL28-mediated RGI (Extended Data Fig. 3a, Supplementary Table 7, Methods 6). Finally, we tested whether the RGI-suppressor strains maintain their capacity to attenuate RGI in the context of the full 185-member community (Methods 7). We compared the root phenotype of seedlings exposed to either the full SynCom or to the same community dropping-out all ten *Variovorax* strains and/or all six *Burkholderia* strains present in this SynCom (drop-out system, Fig. 3a, b, c). We found that *Variovorax* are necessary and sufficient to revert RGI within the full community (Fig. 3b, c, Supplementary Table 8). Further, the presence of *Variovorax* increases the plant’s total root network length (Extended Data Fig. 3b). This result was robust across a range of substrates, including soil, and under various biotic and abiotic contexts (Fig. 3d, e, f, Extended Data Fig. 3b, 4, Supplementary Table 8; Methods 8, 9).

**Fig. 3.**
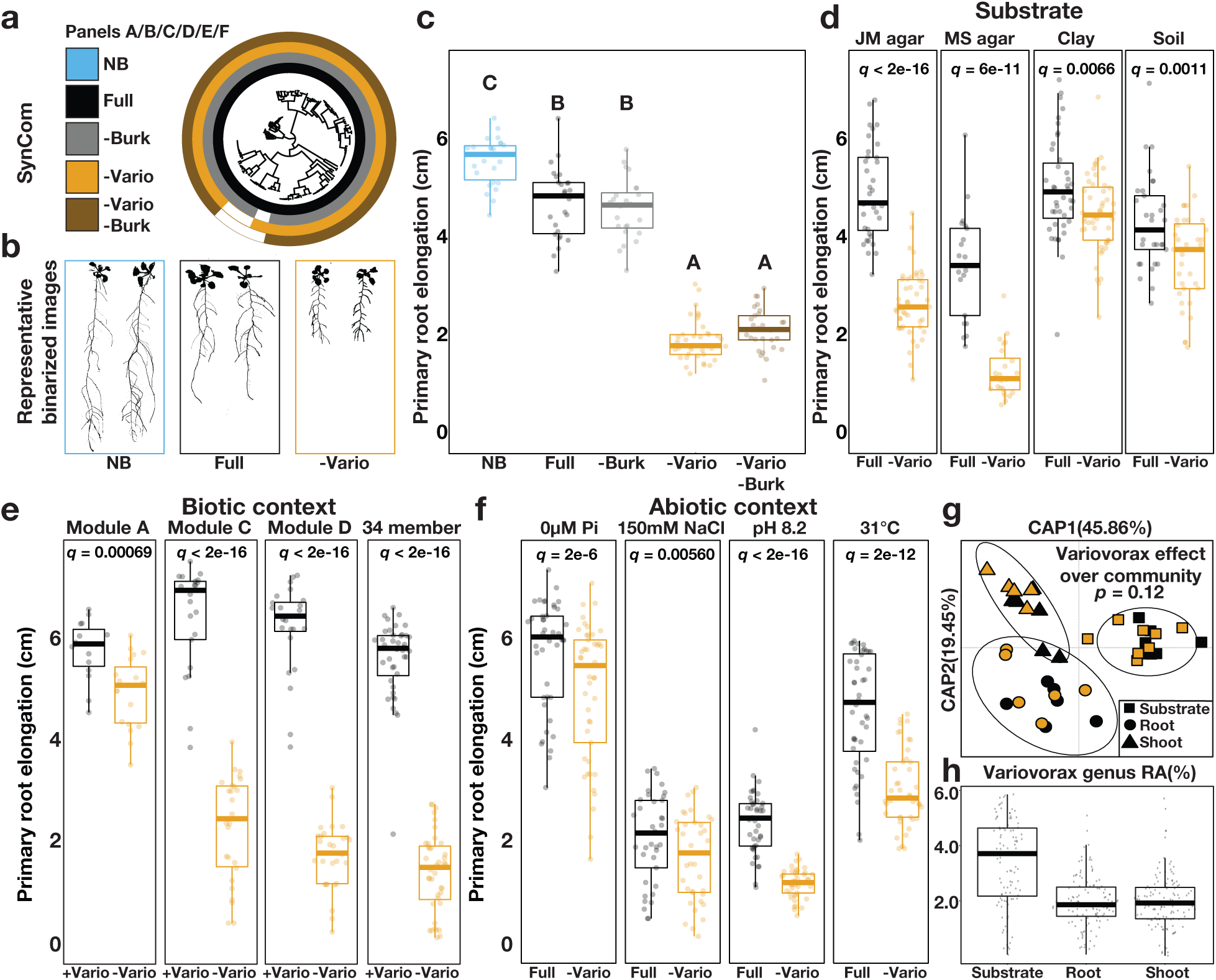
Variovorax are necessary and sufficient to maintain stereotypic root development. **(a)** Phylogenetic tree of 185 bacterial isolates. Concentric rings represent isolate composition of each SynCom treatment (-Burk: Burkholderia drop-out, -Vario: Variovorax drop-out). **(b)** Binarized image of representative uninoculated seedlings (NB), or seedlings with the full SynCom (Full) or the Variovorax drop-out SynCom (-Vario) treatments. **(c)** Primary root elongation of uninoculated seedlings (NB) or seedlings with the different SynCom treatments. Letters indicate statistical significance. **(d)** Primary root elongation of seedlings inoculated with the Full SynCom or with the Variovorax drop-out SynCom (-Vario) across different substrates: Johnson Medium (JM agar), Murashige and Skoog (MS agar), or pots with sterilized clay or potting soil. **(e)** Primary root elongation of seedlings inoculated independently with four compositionally different SynComs (Module A, C, D and 34-member) with (Full) or without (Vario) 10 Variovorax isolates. **(f)** Primary root elongation of seedlings inoculated with the Full SynCom or with the Variovorax drop-out SynCom (-Vario) across different abiotic conditions: unamended medium (JM agar control), phosphate starvation (JM agar 0 µM Pi), salt stress (JM agar 150 mM NaCl), high pH (JM agar pH 8.2) and high temperature (JM agar 31°C). FDR-corrected p-values are shown within each plot. **(g)** Canonical analysis of principal coordinates scatterplots comparing community full vs Variovorax drop-out SynComs across all fractions (agar, root, shoot). PERMANOVA p-value is shown. **(h)** Relative abundance (RA) of the Variovorax genus within the full SynCom across the agar, root and shoot fractions.

To ascertain the phylogenetic breadth of the *Variovorax* ability to attenuate RGI, we tested additional *Variovorax* strains from across the genus’ phylogeny (Extended Data Fig. 5, Supplementary Table 1; Methods 10). All 19 tested *Variovorax* reverted RGI induced by *Arthrobacter* CL28 (Methods 11). A strain from the nearest plant-associated outgroup to this genus, *Acidovorax* Root219^26^, did not revert RGI (Extended Data Fig. 5, Supplementary Table 9). Thus, all tested strains representing the broad phylogeny of *Variovorax* interact with a wide diversity of bacteria to enforce stereotypic root development within complex communities, independent of biotic or abiotic contexts. We tested whether *Variovorax* attenuate RGI by inhibiting growth of RGI-inducing strains (Methods 12). We counted colony forming units of the RGI inducer *Arthrobacter* CL28 from roots in the presence or absence of *Variovorax* CL14 and found that CL28 abundance increased in the presence of *Variovorax* CL14 (Extended Data Fig. 6, Supplementary Table 10). To test whether *Variovorax* modulates bacterial abundances in the whole community, we compared the bacterial relative abundance profiles in seedlings colonized with the full SynCom to that colonized with the *Variovorax* drop-out community (Methods 8d). We found no changes in the abundances of RGI-inducing strains in response to the *Variovorax* drop-out (Fig. 3g). Notably, *Variovorax* account for only ∼1.5% of the root community (Fig. 3h). These results rule out the possibility that *Variovorax* enforce stereotypic root developmentby antagonizing or outcompeting RGI-inducers.

### *Variovorax* maintain root development through auxin and ethylene manipulation

To study the mechanisms underlying bacterial effects on root development, we analysed the transcriptomes of seedlings colonized for 12 days with the RGI-inducing strain *Arthrobacter* CL28 and the RGI-attenuator strain *Variovorax* CL14, either in mono-association with the seedling or in a tripartite combination (Fig. 2f; Methods 13). We also performed RNA-Seq on seedlings colonized with the full SynCom (no RGI) or the *Variovorax* drop-out SynCom (RGI; Fig. 3a). Eighteen genes were significantly induced only under RGI conditions across both experiments (Fig. 4a, b, Supplementary Table 11). Seventeen of these are co-expressed genes with proposed functions related to the root apex^27^ (Fig 4b, Extended Data Fig. 7). The remaining gene is indole-3-acetic acid-amido synthetase GH3.2, which conjugates excess amounts of the plant hormone auxin and is a robust marker for late auxin responses^28, 29^ (Fig. 4b). Production of auxins is a well-documented mechanism by which bacteria modulate plant root development^13^. Indeed, the top 12 auxin-responsive genes from an RNA-Seq study examining acute auxin response in Arabidopsis^28^ exhibited an average transcript increase in seedlings exposed to our RGI-inducing conditions (Fig. 4c and Supplementary Table 12). We hypothesized that RGI attenuation by *Variovorax* is likely mediated by interference with bacterially-produced auxin signalling.

**Fig. 4.**
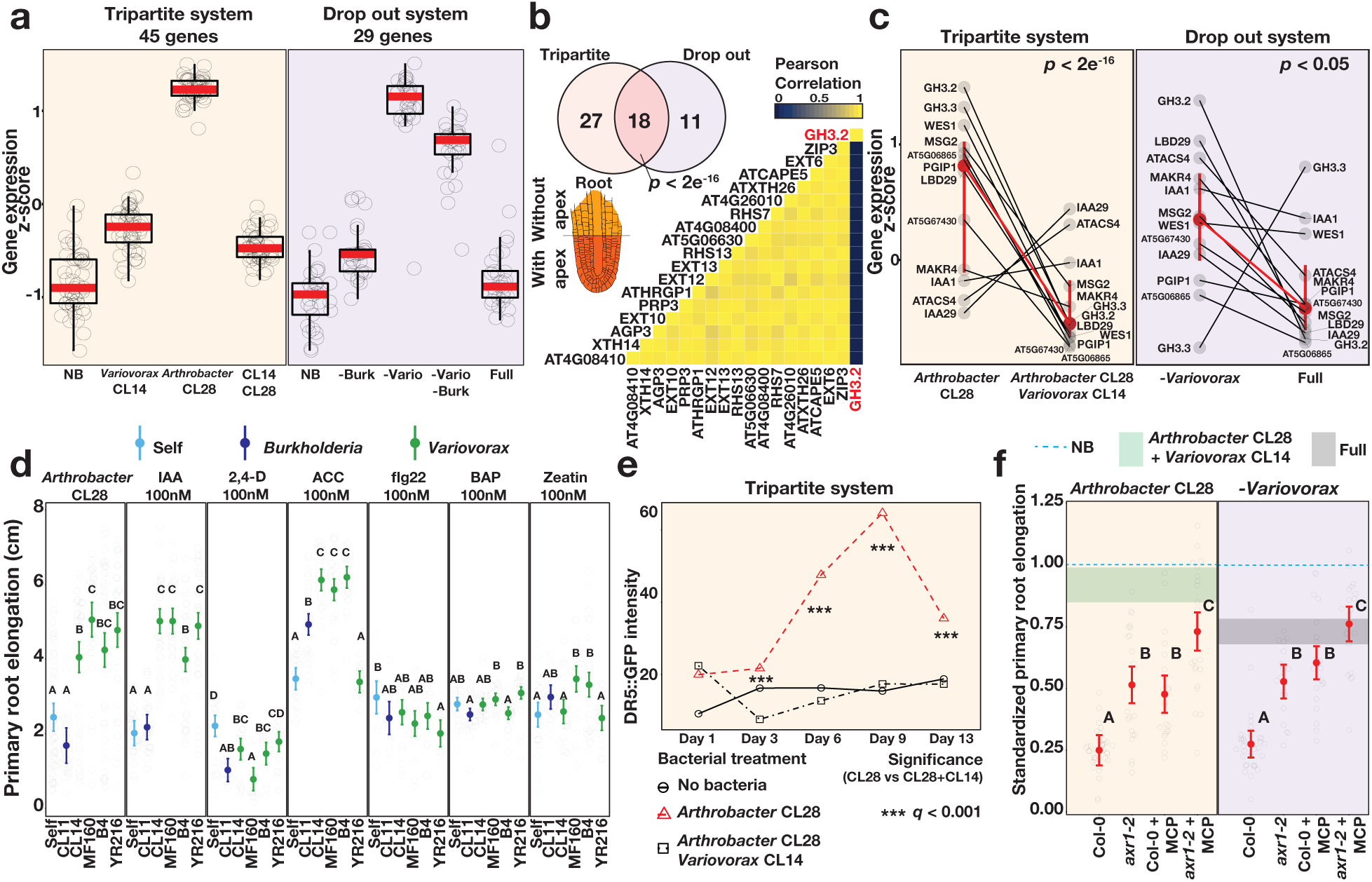
Variovorax attenuation of root growth inhibition is related to auxin and ethylene signaling. **(a)** Boxplots showing the average standardized expression of genes induced in seedlings in response to: Left (Tripartite system) Arthrobacter CL28 compared with uninoculated seedlings (NB) or seedlings inoculated with both Arthrobacter CL28 and Variovorax CL14 (CL14 CL28). Right (Drop-out System) Variovorax drop-out SynCom (-Vario) compared to uninoculated seedlings (NB) and to the full SynCom (Full). **(b)** Venn diagram showing the overlap of enriched genes between the tripartite and drop-out systems. The heatmap shows the pairwise correlation in expression of these 18 genes across tissues27. **(c)** Standardized expression of 12 late-responsive auxin genes across the tripartite and drop-out systems. Each dot represents a gene. Identical genes are connected between bacterial treatments with a black line. Mean expression (95% CI intervals) of the aggregated 12 genes in each treatment is highlighted in red and connected between bacterial treatments with a red line. **(d)** Primary root elongation of seedlings grown with six hormone or MAMP RGI treatments (panels) individually (Self) or with either Burkholderia CL11 or four Variovorax isolates. Significance between the bacterial treatments is shown using the confidence letter display. **(e)** GFP intensity of DR5::GFP Arabidopsis seedlings grown with no bacteria, Arthrobacter CL28 and Arthrobacter CL28+Variovorax CL14. Significance within time points is denoted with asterisks. **(f)** Primary root elongation, standardized to sterile conditions, of wild type (Col-0) auxin unresponsive (axr1-2), ethylene unresponsive (Col-0 + MCP), or auxin/ethylene unresponsive (axr1-2 + MCP) seedlings inoculated with RGI-inducing Arthrobacter CL28 or the Variovorax dropout SynCom (-Variovorax). The blue dotted line marks the relative mean length of uninoculated seedlings. The horizontal shade in each panel corresponds to the interquartile range of seedlings grown with: Arthrobacter CL28+Variovorax CL14), or the full 185-member SynCom including 10 Variovorax isolates (Full SynCom). Differences between treatments are denoted using the compact letter display.

We asked whether RGI-attenuation by *Variovorax* is directly and exclusively related to auxin signalling (Methods 14). Besides auxin, other small molecules cause RGI. These include the plant hormones ethylene^30^ and cytokinin^31^; and microbial-associated molecular patterns (MAMPs) including the flagellin-derived peptide flg22^32^. We tested the ability of diverse *Variovorax* strains and of the *Burkholderia* strain CL11 to revert RGI induced by auxins (Indole-3-acetic acid [IAA] and the auxin analogue 2,4-Dichlorophenoxyacetic acid [2,4-D]), ethylene (the ethylene precursor 1-Aminocyclopropane-1-carboxylic acid [ACC]), cytokinins (Zeatin, 6-Benzylaminopurine) and flg22 peptide (Fig. 4d). All tested *Variovorax* suppressed RGI induced by IAA or ACC (Fig. 4d, Supplementary Table 13), with the exception of *Variovorax* YR216 which did not suppress ACC-induced RGI and does not contain an ACC deaminase gene (Extended Data Fig. 5a), a plant growth-promoting feature associated with this genus^30^. *Burkholderia* CL11 only partially reverted ACC-induced RGI. None of the *Variovorax* attenuated RGI induced by 2,4-D, by flg22 or by cytokinins, indicating that *Variovorax* revert RGI induction by interfering with auxin and/or ethylene signalling. Furthermore, this function is mediated by recognition of auxin by *Variovorax* and not by the plant auxin response per se, since the auxin response (RGI) induced by 2,4-D is not reverted. Indeed, we found that *Variovorax* CL14 degrades IAA in culture (Extended Data Fig. 8a; Methods 15) and quenches fluorescence of the Arabidopsis auxin reporter line *DR5::GFP* caused by the RGI inducer *Arthrobacter* CL28 (Fig. 4e, Extended Data Fig. 8b, Supplementary Table 14; Methods 17).

To ascertain the roles of auxin and ethylene perception by the plant in responding to RGI-inducers, we used the auxin-insensitive *axr2-1* mutant^33^, combined with a competitive inhibitor of ethylene receptors, 1-Methylcyclopropene (1-MCP)^34^, (Methods 17). We inoculated wild type seedlings and the *axr2-1* mutants, treated or not with 1-MCP, with the RGI-inducing *Arthrobacter* CL28 or the *Variovorax* drop-out SynCom (Supplementary Table 15). We observed in both cases that bacterial RGI is reduced in *axr2-1* and 1-MCP-treated wild type seedlings, and is further reduced in doubly-insensitive 1-MCP-treated *axr2-1* seedlings, demonstrating an additive effect of auxin and ethylene on bacterially-induced RGI (Fig. 4f). Thus, a complex SynCom can induce severe morphological changes in root phenotypes via both auxin- and ethylene-dependent pathways, and both are reverted when *Variovorax* are present.

### An auxin catabolism operon in *Variovorax* is required for maintenance of stereotypic plant root development

To identify the bacterial mechanisms involved in RGI-attenuation, we compared the genomes of the 10 *Variovorax* strains in the SynCom to the genomes of the other 175 SynCom members. We identified a list of 947 orthogroups that were rare (< 5% prevalence) in the 175 SynCom members and core (100% prevalence) across the 10 *Variovorax* strains. We collapsed these orthogroups into hotspots (regions of physically continuous orthogroups) and focused on the 12 hotspots that contained at least 10 *Variovorax*-unique orthogroups (Extended Data Fig. 9a, Methods 18 and Supplementary Table 16). Hotspot 33 contains genes with low sequence homology (average ∼30 % identity) to the genes *iacCDEFR* of the *Paraburkholderia phytofirmans* PsJN IAA-degradation *iac* operon^19^ but lacks *iacABI* which were shown to be necessary for *Paraburkholderia* growth on IAA^18^ (Fig. 5a). To test whether the hotspots we identified are responsive to RGI-inducing bacteria, we analysed the transcriptome of *Variovorax* CL14 in monoculture and in co-culture with the RGI-inducing strain *Arthrobacter* CL28 (Methods 19). We observed extensive transcriptional reprograming of *Variovorax* CL14 when co-cultured with *Arthrobacter* CL28 (Supplementary Table 17). Among the 12 hotspots we identified, the genes in hotspot 33 were the most highly upregulated (Fig. 5a, Extended Data Fig. 9b, Methods 19). We thus hypothesized that hotspot 33 contains an uncharacterized auxin degradation operon.

**Fig. 5.**
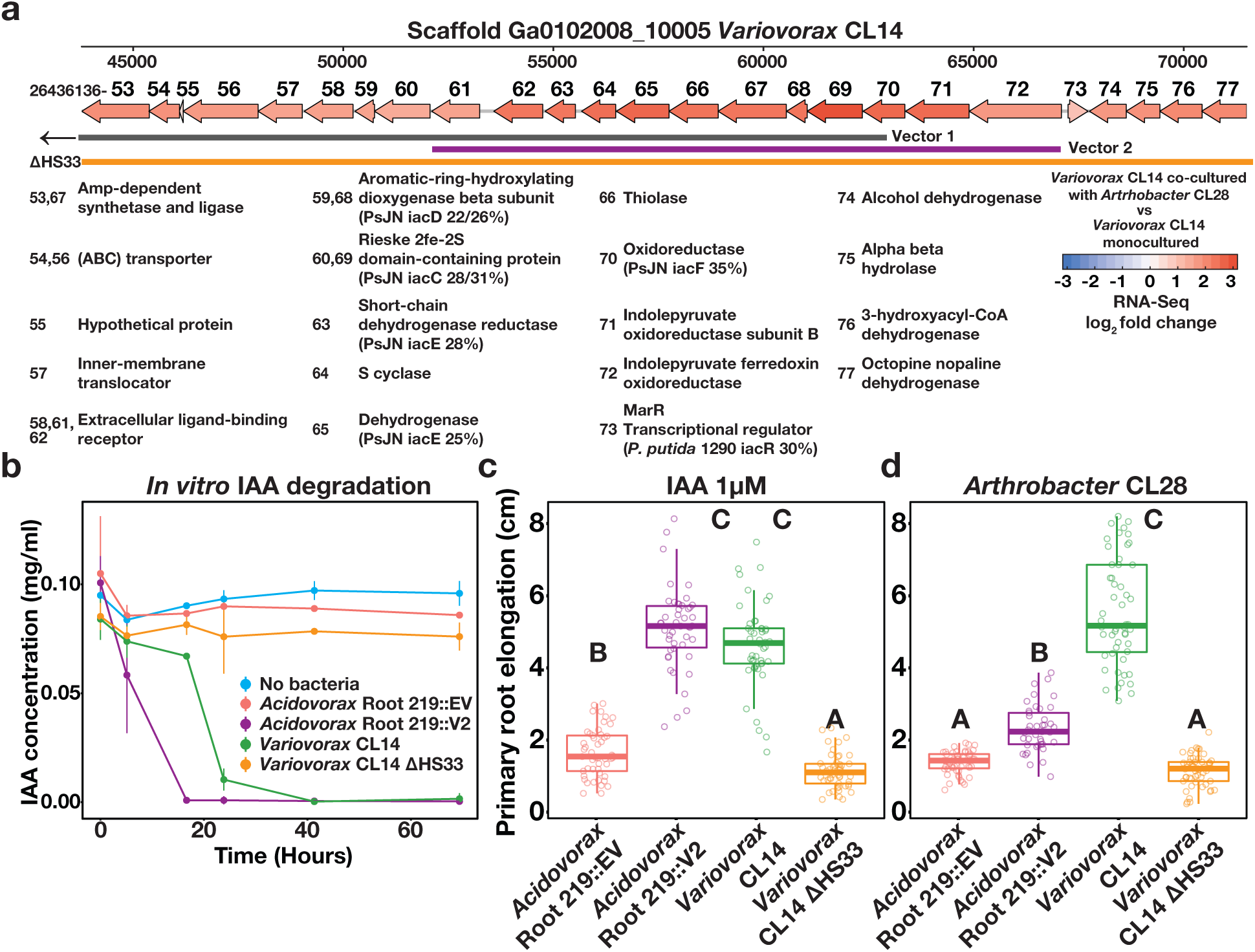
An auxin-degrading operon in Variovorax is required for maintenance of stereotypical root development. **(a)** A map of the hotspot 33. Genes are annotated with the last two digits of their IMG gene ID (26436136XX) and their functional assignments are shown below the map, including % identity of any to genes from a known auxin degradation locus. Gene are colored by the log2 fold change in their transcript abundance in Variovorax CL14 co-cultured with Arthrobacter CL28 vs Variovorax CL14 monoculture. The overlap of this region vectors 1 and 2 and the region knocked out in Variovorax CL14L1HS33 are shown below the map. Note that vector 1 extends beyond this region. **(b)** In-vitro degradation of IAA by Acidovorax 219::EV, Acidovorax 219::V2, Variovorax CL14 and Variovorax CL14L1HS33. **(c)** Primary root elongation of seedlings treated with IAA alone or inoculated with Acidovorax 219::EV, Acidovorax 219::V2, Variovorax CL14 and Variovorax CL14L1HS33. **(d)** Primary root elongation of seedlings inoculated with Arthrobacter CL28 in alone or together with Acidovorax 219::EV, Acidovorax 219::V2, Variorovax CL14, or Variovorax CL14 L1HS33. Letters indicate post-hoc significance.

In parallel, we constructed a *Variovorax* CL14 genomic library in *E. coli* with >12.5 kb inserts in the broad host range vector *pBBR1-MCS2*, and screened the resulting *E. coli* clones for auxin degradation (Methods 20). Two clones from the approximately 3,500 screened degraded IAA (Fig. 5a Vector 1 and Vector 2, Fig. 5b, Methods 20b). The *Variovorax* CL14 genomic inserts in both of these clones contained portions of hotspot 33 (Fig. 5a, Extended Data Fig. 9). The overlap common to these clones contained 9 genes, among them the weak homologs to *Paraburkholedria iacCDE*. To test whether this genomic region is sufficient to revert RGI in the plant, we transformed a relative of *Variovorax*, *Acidovorax* Root219 that does not cause or revert RGI (Extended Data Fig. 5), with the shorter functional insert containing Vector 2 (V2) or with an empty vector (EV). We inoculated the resulting gain-of-function strain, *Acidovorax* Root219::V2 or the control *Acidovorax* Root219::EV onto plants treated with IAA or inoculated with the RGI strain *Arthrobacter* CL28. *Acidovorax* Root219::V2 fully reverted IAA-induced RGI (Fig. 5c) and partially reverted *Arthrobacter* CL28-induced RGI, despite colonizing roots at significantly lower numbers than *Variovorax* CL14 (Fig.5d, Supplementary table 18). Critically, we deleted hotspot 33 from *Variovorax* CL14 (Fig. 5a) to test whether this putative operon is necessary for RGI-reversion. The resulting strain *Variovorax* CL14ΔHS33 did not degrade IAA in culture (Fig. 5b) and did not revert IAA-induced (Fig. 5c) or *Arthrobacter* CL28-induced (Fig. 5d) RGI. Thus, this *Variovorax*-specific gene cluster is necessary for the RGI-suppression phenotype and auxin degradation. It is thus the critical genetic locus required by *Variovorax* to maintain stereotypic root development in the context of a phylogenetically diverse microbiome.

## Conclusions

Signalling molecules and other secondary metabolites are adaptations that allow microbes to survive competition for primary metabolites. Our results illuminate the importance of a trophic layer of microbes that utilize these secondary metabolites for their own benefit, while potentially providing the unselected exaptation^35^ of interfering with signalling between the bacterial microbiota and the plant host. Metabolic signal interference was demonstrated in the case of quorum quenching^15^, degradation of MAMPs^36^ and, indeed, the degradation of bacterially-produced auxin, including among *Variovorax*^13, 14, 16^. We have shown here that the metabolic homeostasis enforced by the presence of *Variovorax* in a phylogenetically diverse, realistic synthetic community allows the plant to maintain its developmental program within a chemically complex matrix. *Variovorax* were recently found to have the rare property of improved plant-colonization upon late arrival to an established community, as opposed to arriving together with the other strains in the community^37^, suggesting that *Variovorax* utilize bacterially-produced/induced rather than plant-derived substrates. Furthermore, via re-analysis of a recent large-scale time- and spatially-resolved survey of the Arabidopsis root microbiome^38^, we noted that two *Variovorax*-16S sequences were found in ∼85% and 100% of the sampled sites (among the top 10 most prevalent sequence among rhizoplane samples, Extended Data Fig. 10). These ecological observations, together with our results using a reductionist microcosm, reinforce the importance of *Variovorax* as key players in bacteria-bacteria-plant communication networks required to maintain stereotypic root development within a complex biochemical ecosystem.

## Methods

### 1. Arabidopsis with bacterial SynCom microcosm across four stress gradients (Fig. 1, fig. S2-S3, data S2)

#### a. Bacterial culture and plant-inoculation

The 185-member bacterial synthetic community (SynCom) used here contains genome-sequenced isolates obtained from surface sterilized Brassicaceae roots, nearly all *Arabidopsis thaliana*, planted in two North Carolina, US, soils. A detailed description of this collection and isolation procedures can be found in^24^. One week prior to each experiment, bacteria were inoculated from glycerol stocks into 600 µL KB medium in a 96 deep well plate. Bacterial cultures were grown at 28 °C, shaking at 250 rpm. After five days of growth, cultures were inoculated into fresh media and returned to the incubator for an additional 48 hours, resulting in two copies of each culture, 7 days old and 48 hours old. We adopted this procedure to account for variable growth rates of different SynCom members and to ensure that non-stationary cells from each strain were included in the inoculum. After growth, 48-hour and 7-day plates were combined and optical density of cultures was measured at 600 nm (OD_600_) using an Infinite M200 Pro plate reader (TECAN). All cultures were then pooled while normalizing the volume of each culture to OD_600_=1. The mixed culture was washed twice with 10 mM MgCl2 to remove spent media and cell debris and vortexed vigorously with sterile glass beads to break up aggregates. OD_600_ of the mixed, washed culture was then measured and normalized to OD_600_=0.2. The SynCom inoculum (100 µL) was spread on 12 X 12 cm vertical square agar plates with amended Johnson medium (JM)^2^ without sucrose prior to transferring seedlings.

#### b. In vitro plant growth conditions

All seeds were surface-sterilized with 70% bleach, 0.2% Tween-20 for 8 min, and rinsed three times with sterile distilled water to eliminate any seed-borne microbes on the seed surface. Seeds were stratified at 4 °C in the dark for two days. Plants were germinated on vertical square 12 X 12 cm agar plates with JM containing 0.5% sucrose, for 7 days. Then, 10 plants were transferred to each of the SynCom-inoculated agar plates. The composition of JM in the agar plates was amended to produce environmental variation. We added to the previously reported phosphate concentration gradient (0, 10, 30, 50, 100, 1000 µM Pi)^39^ three additional environmental gradients: Salinity (50, 100, 150, 200 mM NaCl), pH (5.5, 7.0, 8.2), and incubation temperature (10, 21, 31°C). Each gradient was tested separately, in two independent replicas. Each condition included three SynCom+plant samples, two no plant controls and one no bacteria control. Plates were placed in randomized order in growth chambers and grown under a 16-h dark/8-h light regime at 21 °C day/18 °C night for 12 days. Upon harvest, DNA was extracted from roots, shoots and agar.

#### c. DNA extraction

Roots, shoots and agar were harvested separately, pooling 6-8 plants for each sample. Roots and shoots were placed in 2 mL Eppendorf tubes with three sterile glass beads. These samples were washed three times with sterile distilled water to remove agar particles and weakly associated microbes. Tubes containing the samples were stored at -80 °C until processing. Root and shoot samples were lyophilized for 48 hours using a Labconco freeze dry system and pulverized using a tissue homogenizer (MPBio). Agar from each plate was collected in 30 mL syringes with a square of sterilized Miracloth (Millipore) at the bottom and kept at -20 °C for one week. Syringes were then thawed at room temperature and samples were squeezed gently through the Miracloth into 50 mL falcon tubes. Samples were centrifuged at max speed for 20 min and most of the supernatant was discarded. The remaining 1-2 mL of supernatant, containing the pellet, was transferred into clean 1.5 mL Eppendorf tubes. Samples were centrifuged again, supernatant was removed, and pellets were stored at -80 °C until DNA extraction. DNA extractions were carried out on ground root and shoot tissue and agar pellets using 96-well-format MoBio PowerSoil Kit (MOBIO Laboratories; Qiagen) following the manufacturer’s instruction. Sample position in the DNA extraction plates was randomized, and this randomized distribution was maintained throughout library preparation and sequencing.

#### d. Bacterial 16S sequencing

We amplified the V3-V4 regions of the bacterial 16S rRNA gene using the primers 338F (5′-ACTCCTACGGGAGGCAGCA-3′) and 806R (5′-GGACTACHVGGGTWTCTAAT-3′). Two barcodes and six frameshifts were added to the 5’ end of 338F and six frameshifts were added to the 806R primers, based on the protocol by Lundberg et al^40^. Each PCR reaction was performed in triplicate, and included a unique mixture of three frameshifted primer combinations for each plate. PCR conditions were as follows: 5 μL Kapa Enhancer, 5 μL Kapa Buffer A, 1.25 μL 5 μM 338F, 1.25 μL 5 μM 806R, 0.375 μL mixed plant rRNA gene-blocking peptide nucleic acids (PNAs; 1:1 mix of 100 μM plastid PNA and 100 μM mitochondrial PNA^40^), 0.5 μL Kapa dNTPs, 0.2 μL Kapa Robust Taq, 8 μL dH_2_O, 5 μL DNA; temperature cycling: 95°C for 60 s; 24 cycles of 95°C for 15 s; 78°C (PNA) for 10 s; 50°C for 30 s; 72°C for 30 s; 4°C until use. Following PCR cleanup, using AMPure beads (Beckman Coulter), the PCR product was indexed using 96 indexed 806R primers with the Kapa HiFi Hotstart readymix with the same primers as above; temperature cycling: 95°C for 60 s; 9 cycles of 95°C for 15 s; 78°C (PNA) for 10 s; 60°C for 30 s; 72°C for 35 s; 4°C until use. PCR products were purified using AMPure XP magnetic beads (Beckman Coulter) and quantified with a Qubit 2.0 fluorometer (Invitrogen). Amplicons were pooled in equal amounts and then diluted to 10 pM for sequencing. Sequencing was performed on an Illumina MiSeq instrument using a 600-cycle V3 chemistry kit. DNA sequence data for this experiment is available at the NCBI bioproject repository (accession PRJNA543313). The abundance matrix, metadata and taxonomy are available at https://github.com/isaisg/variovoraxRGI.

#### e. 16S amplicon sequence data processing

SynCom sequencing data were processed with MT-Toolbox^41^. Usable read output from MT-Toolbox (that is, reads with 100% correct primer and primer sequences that successfully merged with their pair) were quality filtered using Sickle^42^ by not allowing any window with Q-score under 20. The resulting sequences were globally aligned to a reference set of 16S rDNA sequences extracted from genome assemblies of SynCom members. For strains that did not have an intact 16S rDNA sequence in their assembly, we sequenced the 16S rRNA gene using Sanger sequencing. The reference database also included sequences from known bacterial contaminants and Arabidopsis organellar sequences. Sequence alignment was performed with USEARCH v7.1090^43^ with the option ‘usearch_global’ at a 98% identity threshold. On average, 85% of sequences matched an expected isolate. Our 185 isolates could not all be distinguished from each other based on the V3-V4 sequence and were thus classified into 97 unique sequences (USeqs). A USeq encompasses a set of identical (clustered at 100%) V3-V4 sequences coming from a single or multiple isolates.

Sequence mapping results were used to produce an isolate abundance table. The remaining unmapped sequences were clustered into Operational Taxonomic Units (OTUs) using UPARSE^44^ implemented with USEARCH v7.1090, at 97% identity. Representative OTU sequences were taxonomically annotated with the RDP classifier^45^ trained on the Greengenes database^46^ (4 February 2011). Matches to Arabidopsis organelles were discarded. The vast majority of the remaining unassigned OTUs belonged to the same families as isolates in the SynCom. We combined the assigned USeq and unassigned OTU count tables into a single count table. In addition to the raw count table, we created rarefied (1000 reads per sample) and relative abundance versions of the abundance matrix for further analyses.

The resulting abundance tables were processed and analyzed with functions from the ohchibi package (https://github.com/isaisg/ohchibi). An alpha diversity metric (Shannon diversity) was calculated using the diversity function from the vegan package v2.5-3^47^. We used ANOVA to test for differences in alpha diversity between groups. Beta diversity analyses (Principal coordinate analysis, and canonical analysis of principal coordinates) were based on Bray-Curtis dissimilarity calculated from the relative abundance matrices. We used the capscale function from the vegan R package v.2.5-3^47^ to compute the canonical analysis of principal coordinates (CAP). To analyze the full dataset (all fraction, all abiotic treatments), we constrained by fraction and abiotic treatment while conditioning for the replica and experiment effect. We explored the abiotic conditions effect inside each of the four abiotic gradients tested (phosphate, salinity, pH and temperature). We performed the Fraction:abiotic interaction analysis within each fraction independently, constraining for the abiotic conditions while conditioning for the replica effect. In addition to CAP, we performed Permutational Multivariate Analysis of Variance (PERMANOVA) using the adonis function from the vegan package v2.5-3^47^. We used the package DESeq2 v1.22.1^48^ to compute the enrichment profiles for USeqs present in the count table.

We estimated the fraction effect across all the abiotic conditions tested by creating a group variable that merged the fraction variable and the abiotic condition variable together (e.g Root_0Pi, Agar_0Pi). We fitted the following model specification using this group variable:

Abundance ∼ Rep + Experiment + group

From the fitted model, we extracted, for all levels within the group variables, the following comparisons: Agar vs Root and Agar vs Shoot. A USeq was considered statistically significant if it had a false discovery rate (FDR) adjusted *p*-value < 0.05.

All scripts and dataset objects necessary to reproduce the synthetic community analyses are deposited in the following github repository: https://github.com/isaisg/variovoraxRGI

#### f. Co-occurrence analysis

The relative abundance matrix (USeqs X Samples) was standardized across the USeqs by dividing the abundance of each USeq in its sample over the mean abundance of that USeq across all samples. Subsequently, we created a dissimilarity matrix based on the Pearson correlation coefficient between all the pairs of strains in the transformed abundance matrix, using the cor function in the stats base package in R. Finally, hierarchichal clustering (method ward.D2, function hclust) was applied over the dissimilarity matrix constructed above.

#### g. Heatmap and family enrichment analysis

We visualized the results of the GLM model testing the fraction effects across each specific abiotic condition tested using a heatmap. The rows in the heatmap were ordered according to the dendrogram order obtained from the USeqs co-occurrence analysis. The heatmap was colored based on the log2FoldChange output by the GLM model. We highlighted in a black shade the comparisons that were significant (*q*-value < 0.05). Finally, for each of the four modules we computed for each family present in that module a hypergeometric test testing if that family was overrepresented (enriched) in that particular module. Families whose FDR *p*-value < 0.1 were visualized in the figure.

### 2. Deconstructing the SynCom to four modules of co-occurring strains (Fig. 2a, 2c and data S3)

#### a. Bacterial culture and plant-inoculation

Strains belonging to each module: A, B, C and D (Materials and Methods 1f) were grown in separate deep 96-well plates and mixed as described above (Materials and Methods 1a). The concentration of each module was adjusted to OD_600_=0.05 (1/4 of the concentration of the full SynCom). Each module was spread on the plates either separately, or in combination with another module at a total volume of 100 µL. In addition, we included a full SynCom control and an uninoculated control, bringing the number of SynCom combinations to 12. We performed the experiment in two independent replicates and each replicate included five plates per SynCom combination.

#### b. *In vitro* plant growth conditions

Seed sterilization and germination conditions were the same as Materials and Methods 1b. Plants were transferred to each of the SynCom-inoculated agar plates containing JM without sucrose. Plates were placed in randomized order in growth chambers and grown under a 16-h dark/8-h light regime at 21 °C day/18 °C night for 12 days. Upon harvest, root morphology was measured.

#### c. Root and shoot image analysis

Plates were imaged twelve days post-transferring, using a document scanner. Primary root length elongation was measured using ImageJ^49^ and shoot area and total root network were measured with WinRhizo software (Regent Instruments Inc.).

#### d. Primary root elongation analyses

Primary root elongation was compared across the No Bacteria, full SynCom, single modules and pairs of modules treatments jointly using an ANOVA model controlling for the replicate effect. Differences between treatments were indicated using the confidence letter display (CLD) derived from the Tukey’s post hoc test implemented in the package emmeans^50^.

### 3. Inoculating plants with all SynCom isolates separately (Fig. 2B, fig. S4 and data S4)

#### a. Bacterial culture and plant-inoculation

Cultures from each strain in the SynCom were grown in KB medium and washed separately (Methods 1a), and OD_600_ was adjusted to 0.01 before spreading 100 µL on plates. We performed the experiment in two independent replicates and each replicate included one plate per each of the 185 strains. *In vitro* growth conditions were the same as in Materials and Methods 2b. Upon harvest, root morphology was measured (Materials and Methods 2c). Isolates generating an average main root elongation of <3 cm were classified as RGI-inducing strains.

### 4. Tripartite plant-microbe-microbe experiments (Fig. 2D-F and data S5-S6)

#### a. Experimental design

To identify strains that revert RGI (Fig. 2D and data S5), we selected all 18 non-RGI inducing strains in module A and co-inoculated them with each of four RGI inducing strains, one from each module. The experiment also included uninoculated controls and controls consisting of each of the 22 strains inoculated alone, amounting to 95 separate bacterial combinations.

To confirm the ability of *Variovorax* and *Burkholderia* to attenuate RGI induced by diverse bacteria (Fig. 2E and data S6), three RGI attenuating strains were co-inoculated with a selection of 18 RGI inducing strains. The experiment also included uninoculated controls and controls consisting of each of the 21 strains inoculated alone. Thus, the experiment consisted of 76 separate bacterial combinations. We performed each of these two experiments in two independent replicates and each replicate included one plate per each of the strain combinations.

#### b. Bacterial culture and plant-inoculation

All strains were streaked on agar plates, then transferred to 4 ml liquid KB medium for over-night growth. Cultures were then washed, and OD_600_ was adjusted to 0.02 before mixing and spreading 100 µL on each plate. Upon harvest, root morphology was measured (Materials and Methods 2c) and plant RNA was harvested and processed from uninoculated samples, and from samples with *Variovorax* CL14, *Arthrobacter* CL28 and the combination of both (Materials and Methods 4d).

#### c. Primary root elongation analysis

We fitted ANOVA models for each RGI-inducing strain tested. Each model compared the primary root elongation with the RGI inducing strains alone against root elongation when the RGI inducing strain was co-inoculated with other isolates. The *p*-values for all the comparisons were corrected for multiple testing using false discovery rate (FDR).

#### d. RNA extraction

RNA was extracted from *A. thaliana* seedlings following Logemann et al^51^. Four seedlings were harvested from each sample and samples were flash frozen and stored at -80 °C until processing. Frozen seedlings were ground using a TissueLyzer II (Qiagen), then homogenized in a buffer containing 400 μL of Z6-buffer; 8 M guanidine HCl, 20 mM MES, 20 mM EDTA at pH 7.0. 400 μL phenol:chloroform:isoamylalcohol, 25:24:1 was added, and samples were vortexed and centrifuged (20,000 g, 10 minutes) for phase separation. The aqueous phase was transferred to a new 1.5 mL Eppendorf tube and 0.05 volumes of 1 N acetic acid and 0.7 volumes 96% ethanol were added. The RNA was precipitated at -20 °C overnight. Following centrifugation (20,000 g, 10 minutes, 4°C), the pellet was washed with 200 μL sodium acetate (pH 5.2) and 70% ethanol. The RNA was dried and dissolved in 30 μL of ultrapure water and stored at -80 °C until use.

#### e. Plant RNA sequencing

Illumina-based mRNA-Seq libraries were prepared from 1 μg RNA following^4^. mRNA was purified from total RNA using Sera-mag oligo(dT) magnetic beads (GE Healthcare Life Sciences) and then fragmented in the presence of divalent cations (Mg^2+^) at 94°C for 6 minutes. The resulting fragmented mRNA was used for first-strand cDNA synthesis using random hexamers and reverse transcriptase, followed by second-strand cDNA synthesis using DNA Polymerase I and RNAseH. Double-stranded cDNA was end-repaired using T4 DNA polymerase, T4 polynucleotide kinase, and Klenow polymerase. The DNA fragments were then adenylated using Klenow exo-polymerase to allow the ligation of Illumina Truseq HT adapters (D501–D508 and D701–D712). All enzymes were purchased from Enzymatics. Following library preparation, quality control and quantification were performed using a 2100 Bioanalyzer instrument (Agilent) and the Quant-iT PicoGreen dsDNA Reagent (Invitrogen), respectively. Libraries were sequenced using Illumina HiSeq4000 sequencers to generate 50-bp single-end reads.

#### f. RNA-Seq read processing

Initial quality assessment of the Illumina RNA-Seq reads was performed using FastQC v0.11.7^52^. Trimmomatic v0.36^53^ was used to identify and discard reads containing the Illumina adaptor sequence. The resulting high-quality reads were then mapped against the TAIR10 Arabidopsis reference genome using HISAT2 v2.1.0^54^ with default parameters. The featureCounts function from the Subread package^55^ was then used to count reads that mapped to each one of the 27,206 nuclear protein-coding genes. Evaluation of the results of each step of the analysis was performed using MultiQC v1.1^56^. Raw sequencing data and read counts are available at the NCBI Gene Expression Omnibus accession number GSE131158.

### 5. *Variovorax* drop-out experiment (Fig. 3A-C and data S7)

#### a. Bacterial culture and plant-inoculation

The entire SynCom, excluding all 10 *Variovorax* isolates and all five *Burkholderia* isolates was grown and prepared as described above (Materials and Methods 1a). The *Variovorax* and *Burkholderia* isolates were grown in separate tubes, washed and added to the rest of the SynCom to a final OD_600_ of 0.001 (the calculated OD_600_ of each individual strain in a 185-Member SynCom at a total of OD_600_ of 0.2), to form the following five mixtures: (i) Full community: all *Variovorax* and *Burkholderia* isolates added to the SynCom; (ii) *Burkholderia* drop-out: only *Variovorax* isolates added to the SynCom; (iii) *Variovorax* drop-out: only *Burkholderia* isolates added to the SynCom; (iv) *Variovorax* and *Burkholderia* drop-out: no isolates added to the SynCom; (v) Uninoculated plants: no SynCom. The experiment consisted of six plates per SynCom mixture, amounting to 30 plates. Upon harvest, root morphology was measured and analyzed (Materials and Methods 1c,4c); and Bacterial DNA (Materials and Methods 1d) and plant RNA (Materials and Methods 4d-e) were harvested and processed.

### 6. *Variovorax* drop-out under varying abiotic contexts (Fig. 3E and data S7)

#### a. Bacterial culture and plant-inoculation

The composition of JM in the agar plates was amended to produce abiotic environmental variation. These amendments included salt stress (150 mM NaCl), low Phosphate (10 µM Phosphate), high pH (pH 8.2) and high temperature (plates incubated at 31 °C), as well as an un-amended JM control. Additionally, we tested a different media (1/2-strength Murashige and Skoog [MS]) and a soil-like substrate. As a soil-like substrate, we used calcined clay (Diamond Pro), prepared as follows: 100 mL of clay was placed in Magenta GA7 jars. The jars were then autoclaved twice. 40 mL of liquid JM was added to the Magenta jars, with the corresponding bacterial mixture spiked into the media at a final OD_600_ of 5E-4. Four 1-week old seedlings were transferred to each vessel, and vessels were covered with Breath-Easy gas permeable sealing membrane (Research Products International) to maintain sterility and gas exchange.

The entire SynCom, excluding all 10 *Variovorax* isolates was grown and prepared as described above (Materials and Methods 1a). The *Variovorax* isolates were grown in separate tubes, washed and added to the rest of the SynCom to a final OD_600_ of 0.001 (the calculated OD_600_ of each individual strain in a 185-Member SynCom at an OD_600_ of 0.2), to form the following five mixtures: (i) Full community: all *Variovorax* isolates added to the SynCom; (ii) *Variovorax* drop-out: no isolates added to the SynCom; (iii) Uninoculated plants: no SynCom.

We inoculated all three SynCom combinations in all seven abiotic treatments, amounting to 21 experimental conditions. We performed the experiment in two independent replicates and each replicate included three plates per experimental conditions, amounting to 63 plates per replicate. Upon harvest, root morphology was measured (Materials and Methods 2c); and Bacterial DNA (Materials and Methods 1c-e) and plant RNA (Materials and Methods 4d-f) were harvested and processed.

#### b. Root image analysis

For agar plates, roots were imaged as described above (Materials and Methods 2c). For calcined clay pots, four weeks post-transferring, pots were inverted, and whole root systems were gently separated from the clay by washing with water. Root systems were spread over an empty petri dish and scanned using a document scanner.

#### c. Primary root elongation and total root network analysis

Primary root elongation was compared between SynCom treatments within each of the different abiotic contexts tested independently. Differences between treatments were indicated using the confidence letter display (CLD) derived from the Tukey’s post hoc test implemented in the package emmeans.

#### d. Bacterial 16S data analysis

To be able to compare shifts in the community composition of samples treated with and without the *Variovorax* genus, we *in silico* removed the 10 *Variovorax* isolates from the count table of samples inoculated with the Full community treatment. We then merged this count table with the count table constructed from samples inoculated without the *Variovorax* genus (*Variovorax* drop-out treatment). Then, we calculated a relative abundance of each USeq across all the samples using the merged count matrix. Finally, we applied Canonical Analysis of Principal Coordinates (CAP) over the merged relative abundance matrix to control for the replica effect. In addition, we utilized the function adonis from the vegan R package to compute a PERMANOVA test over the merged relative abundance matrix and we fitted a model evaluating the fraction and SynCom (presence of *Variovorax*) effects over the assembly of the community.

### 7. *Variovorax* drop-out under varying biotic contexts (Fig. 3D, fig S7 and data S7)

#### a. Bacterial culture and plant-inoculation

Strains belonging to modules A (excluding *Variovorax*), C and D were grown in separate wells in deep 96-well plates and mixed as described above (Materials and Methods 1a). The concentration of each module was adjusted to OD_600_=0.05 (1/4 of the concentration of the full SynCom). The *Variovorax* isolates were grown in separate tubes, washed and added to the rest of the SynCom to a final OD_600_ of 0.001.

In a separate experiment, the 35-member SynCom used by Castrillo et al^2^ was grown, excluding *Variovorax* CL14, to create a taxonomically diverse, *Variovorax*-free subset of the full 185 community. The concentration of this SynCom was adjusted to OD_600_=0.05. The *Variovorax* isolates were grown in separate tubes, washed and added to the rest of the SynCom to a final OD_600_ of 0.001.

These two experiments included the following mixtures (fig S7 and data S7): (i) Module A excluding *Variovorax*; (ii) Module C; (iii) Module D; (iv) Module A including *Variovorax*; (v) Module C + all 10 *Variovorax*; (vi) Module D + all 10 *Variovorax*; (vii) 35-member SynCom excluding *Variovorax*; (viii) 34-member SynCom + all 10 *Variovorax*; (ix) uninoculated control. The experiment with modules A, C and D was performed in two independent experiments, with two plates per treatment in each. The experiment with the 34-member SynCom was performed once, with 5 plates per treatment. Upon harvest, root morphology was measured (Materials and Methods 2c).

#### b. Primary root elongation analysis

We directly compared differences between the full SynCom and *Variovorax* drop-out treatment using a *t*-test and adjusting the *p*-values for multiple testing using false discovery rate.

### 8. Phylogenetic inference of the SynCom and *Variovorax* isolates (Fig 2A, fig. S1A, S4, S7 and S9A-B)

To build the phylogenetic tree of the SynCom isolates, we used the super matrix approach previously described in^24^. We scanned 120 previously defined marker genes across the 185 isolate genomes from the SynCom utilizing the hmmsearch tool from the hmmer v3.1b2^57^. Then, we selected 47 markers that were present as single copy genes in 100% of our isolates. Next, we aligned each individual marker using MAFFT^58^ and filtered low quality columns in the alignment using trimAl^59^. Then, we concatenated all filtered alignments into a super alignment. Finally, FastTree v2.1^60^ was used to infer the phylogeny utilizing the WAG model of evolution. For the *Variovorax* relative’s tree, we chose 56 markers present as single copy across 124 Burkholderiales isolates and implemented the same methodology described above.

### 9. Measuring how prevalent the RGI attenuation trait is across the *Variovorax* phylogeny (fig. S9A-B, data S1 and data S8)

#### a. Bacterial culture and plant-inoculation

Fifteen *Variovorax* strains from across the genus’ phylogeny were each co-inoculated with the RGI inducer *Arthrobacter* CL28. All 16 strains were grown in separate tubes, then washed, and OD_600_ was adjusted to 0.01 before mixing. Pairs of strains were mixed in 1:1 ratios and spread at a total volume of 100 µL onto agar prior to seedling transfer. The experiment also included uninoculated controls and controls consisting of each of the 16 strains inoculated alone. Thus, the experiment consisted of 32 separate bacterial combinations. We performed the experiment one time, which included 3 plates per bacterial combination. Upon harvest, root morphology was measured (Materials and Methods 2c). Primary root elongation was analyzed as described above (Materials and Methods 4c).

### 10. Measuring root growth inhibition in tomato seedlings (fig. S10 and data S9)

#### a. Experimental design

This experiment included the following treatments: (i) No bacteria, (ii) *Arthrobacter* CL28, (iii) *Variovorax* CL14 and (iv) *Arthrobacter* CL28 + *Variovorax* CL14. Each treatment was repeated in three separate agar plates with five tomato seedlings per plate. The experiment was repeated in two independent replicates.

#### b. Bacterial culture and plant-inoculation

All strains were grown in separate tubes, then washed, and OD_600_ was adjusted to 0.01 before mixing and spreading (Methods 3b). 400 µL of each bacterial treatment was spread on 20 X 20 agar plates containing JM agar with no sucrose.

#### c. In vitro plant growth conditions

We used Heinz 1706 seeds. All seeds were soaked in sterile distilled water for 15 min, then surface-sterilized with 70% bleach, 0.2% Tween-20 for 15 min, and rinsed five times with sterile distilled water to eliminate any seed-borne microbes on the seed surface. Seeds were stratified at 4 °C in the dark for two days. Plants were germinated on vertical square 10 X 10 cm agar plates with JM containing 0.5% sucrose, for 7 days. Then, 5 plants were transferred to each of the SynCom-inoculated agar plates. Upon harvest, root morphology was measured. (Materials and Methods 2c).

#### c. Primary root elongation analysis

Differences between treatments were indicated using the confidence letter display (CLD) derived from the Tukey’s post hoc test from an ANOVA model.

### 11. Determination of *Arthrobacter* CL28 colony forming units from roots (fig. S11 and data S10)

Arabidopsis seedlings were inoculated with (i) *Arthrobacter* CL28 alone, (ii) *Arthrobacter* CL28 + *Variovorax* CL14 or (iii) *Arthrobacter* CL28 + *Variovorax* B4, as described above (Material and Methods 4b). Each bacterial treatment included four separate plates, with nine seedlings in each plate. Upon harvest, all seedlings were placed in pre-weighed 2 mL Eppendorf tubes containing three glass beads, three seedlings per tube (producing 12 data points per treatment). Roots were weighed, then homogenized using a bead beater (MP Biomedicals). The resulting suspension was serially diluted, then plated on LB agar plates containing 50 µg/mL of Apramycin and colonies were counting after incubation of 48 hours at 28° C.

### 12. Arabidopsis RNA-Seq analysis (Fig. 4A-C, fig. S13 and data S11-12)

#### a. Detection of RGI-induced genes (Fig. 4A-B)

To measure the transcriptional response of the plant to the different SynCom combinations, we used the R package DESeq2 v.1.22.1^48^. The raw count genes matrixes for the dropout and tripartite experiments were used independently to define differentially expressed genes (DEGs). For the analysis of both experiments we fitted the following model specification:

Abundance Gene ∼ SynCom

From the fitted models we derived the following contrasts to obtain differentially expressed genes (DEGs). A gene was considered differentially expressed if it had a *q*-value < 0.1. For the tripartite system (Materials and Methods 4), we performed the following contrasts: *Arthrobacter* CL28 vs No Bacteria (NB) and *Arthrobacter* CL28 vs *Arthrobacter* CL28 co-inoculated with *Variovorax* CL14. The logic behind these two contrasts was to identify genes that were induced in RGI plants (*Arthrobacter* CL28 vs NB) AND repressed by *Variovorax* CL14. For the dropout system (Materials and Methods 5), we performed the following contrasts, *Variovorax* drop-out vs NB, and *Variovorax* drop-out vs full SynCom. The logic behind these two contrasts was identical to the tripartite system: to identify genes that are associated with the RGI phenotype (*Variovorax* drop-out vs NB contrast) AND repressed when *Variovorax* are present (*Variovorax* drop-out vs full SynCom contrast).

For visualization purposes, we applied a variance stabilizing transformation (DESeq2) to the raw count gene matrix. We then standardized each gene expression (z-score) along the samples. We subset DEGs from this standardized matrix and calculated the mean z-score expression value for each SynCom treatment.

To identify the tissue specific expression profile of the 18 intersecting genes between the tripartite and dropout systems, we downloaded the spatial expression profile of each gene from the Klepikova atlas^27^ using the Bio-analytic resource of plant biology platform. Then, we constructed a spatial expression matrix of the 18 genes and computed pairwise Pearson correlation between all pairs of genes. Finally, we applied hierarchical clustering to this correlation matrix.

#### b. Comparison with acute auxin response dataset (Figure 4C)

We applied the variance stabilizing transformation (DESeq2) to the raw count gene matrix. We then standardized each gene expression (z-score) along the samples. From this matrix, we subsetted 12 genes that in a previous study^28^ exhibited the highest fold change between auxin treated and untreated samples. Finally, we calculated the mean z-score expression value of each of these 12 genes across the SynCom treatments. We estimated the statistical significance of the trend of these 12 genes between a pair of SynCom treatments (Full SynCom vs *Variovorax* drop-out, *Arthrobacter* CL28 vs *Arthrobacter* CL28 plus *Variovorax* CL14) using a permutation approach: we estimated a *p*-value by randomly selecting 12 genes 10000 times from the expression matrix and comparing the mean expression between the two SynCom treatments (e.g Full SynCom vs *Variovorax* drop-out) with the actual mean expression value from the 12 genes reported as robust auxin markers.

### 13. Measuring the ability of *Variovorax* to attenuate RGI induced by small molecules (Figure 4D and data S13) indole-3-acetic acid (IAA), 2,4-Dichlorophenoxyacetic acid (2,4-D), ethylene (the ethylene precursor 1-Aminocyclopropane-1-carboxylic acid [ACC]), cytokinins (Zeatin, 6-Benzylaminopurine) and flagellin 22 peptide (flg22) (Fig. 4d)

#### a. Bacterial culture and plant-inoculation

We embedded each of the following compounds in JM plates: 100 nM Indole acetic acid (IAA, Sigma), 1µM IAA, 100 nM 1-Aminocyclopropane-1-carboxylic acid (ACC, Sigma), 100 nM 2,4-Dichlorophenoxyacetic acid (2,4-d, Sigma), 100 nM flagellin 22 (flg22, PhytoTech labs), 100 nM 6-Benzylaminopurine (BAP, Sigma) and 100 nM Zeatin (Sigma). Plates with each compound were inoculate with one of the *Variovorax* strains CL14, MF160, B4 or YR216 or with *Burkholderia* CL11. These strains were grown in separate tubes, then washed, and OD_600_ was adjusted to 0.01 before spreading 100 µL on plates. In addition, we included uninoculated controls for each compound. We also included unamended JM plates inoculated with the RGI inducer *Arthrobacter* CL28 co-inoculated with each of the *Variovorax*/*Burkholderia* strains or alone. Thus, the experiment included 42 individual treatments. The experiment was repeated twice, with three independent replicates per experiment. Upon harvest, root morphology was measured (Materials and Methods 2c).

#### b. Primary root elongation analysis

Primary root elongation was compared between bacterial treatments within each of root growth inhibition treatments tested. Differences between treatments were estimated as described above (Materials and Methods 4c). We plotted the estimated means with 95% confidence interval of each bacterial treatment across the different RGI treatments.

### 14. *In vitro* growth of *Variovorax* (fig. S14)

*Variovorax* CL14 was grown in 5mL cultures for 40 hours at 28 °C in 1x M9 minimal salts media (Sigma M6030) supplemented with 2 mM MgSO_4_, 0.1 mM CaCl_2_, 10 µM FeSO_4_, and a carbon source: either 15 mM succinate alone, 0.4 mM Indole-3-acetic acid (IAA) with 0.5% Ethanol for IAA solubilization, or both. Optical density at 600 nm and IAA concentrations were measured at six time points. IAA concentrations were measured using the Salkowski method modified from^61^. 100 µL of Salkowski reagent (10 mM FeCl_3_ in 35% perchloric acid) was mixed with 50 µL culture supernatant or IAA standards and color was allowed to develop for 30 min prior to measuring the absorbance at 530nm.

### 15. Measuring plant Auxin response *in-vivo* using a bioreporter line (Fig. 4E-F and data S14)

#### a. Bacterial culture and plant-inoculation

7-day old transgenic Arabidopsis seedlings expressing the *DR5::GFP* reporter construct^62^ were transferred onto plates containing: (i) 100 nM IAA, (ii) *Arthrobacter* CL28, (iii) 100 IAA + *Variovorax* CL14, (iv) *Arthrobacter* CL28 + *Variovorax* CL14, (v) the *Variovorax* drop-out SynCom, (vi) the full SynCom, (vii) uninoculated plates. For treatments ii, iii, Bacterial strains were grown in separate tubes, then washed, and OD_600_ was adjusted to 0.01. For treatment iv, OD-adjusted cultures were mixed in 1:1 ratio and spread onto agar prior to seedling transfer. Cultures for treatments v and vi were prepared as described above (Materials and Methods 6a).

#### b. Fluorescence microscopy

GFP fluorescence in the root elongation zone of 8-10 plants per treatment were visualized using a Nikon Eclipse 80i fluorescence microscope at days 1, 3, 6, 9 and 13 post inoculation. The experiment was performed in two independent replicates.

From each root imaged, 10 random 30 X 30 pixel squares were sampled and average GFP intensity was measured using imageJ^49^. Treatments were compared within each time point using ANOVA tests with Tukey’s post hoc in the R base package emmeans. For visualization purposes we plotted the estimated means of each bacterial across the different timepoints.

### 16. Measuring the dual role of auxin and ethylene perception in SynCom-induced RGI (Fig. 4F and data S15)

#### a. Bacterial culture and plant-inoculation

We transferred four 7-day old wild type seedling and four *axr1-2* seedlings to each plate in this experiment. The plates contained one of five bacterial treatments: (i) *Arthrobacter* CL28, (ii) *Arthrobacter* CL28 + *Variovorax* CL14, (iii) *Variovorax* drop-out SynCom, (iv) Full SynCom, (v) uninoculated, prepared as described above (Materials and Methods 15a) Plates were placed vertically inside sealed 86 X 68 cm Ziploc bags. In one of the bags, we placed an open water container with 80 2.5 gram sachets containing 0.014% 1-MCP (Ethylene Buster, Chrystal International BV). In the second bag we added, as a control, an open water container. Both bags were placed in the growth chamber for 12 days. After 6 days of growth, we added 32 additional sachets to the 1-MCP-treated bag to maintain 1-MCP concentrations in the air. Upon harvest, root morphology was measured (Materials and Methods 2c).

#### b. Primary root elongation analysis

Primary root elongation was standardized to the No bacteria control of each genotype, and compared between genotype/1-MCP treatments within the *Arthrobacter* CL28 treatment and the *Variovorax* drop-out SynCom treatment, independently. Differences between treatments were estimated as described above (Materials and Methods 4c). We plotted the estimated means with 95% CI of each bacterial treatment across the four genotypes. We calculated the interquartile range for the Full and *Arthrobacter* CL28/*Variovorax* CL14 treatments pooling the four genotypes/treatments.

### 17. Preparation of binarized plant images (Fig 2C, 3B and fig. S5-S6)

To present representative root images, we increased contrast and subtracted background in imageJ, then cropped the image to select representative roots. Neighboring roots were manually erased from the cropped image.

### 18. Mining *Variovorax* genomes for auxin degradation operons and ACC-deaminase genes

We used local alignment (BLASTp) to search for the presence of the 10 genes (*iacABCDEFGHIY*) from a previously characterized auxin degradation operon in a different genus^18^ across the 10 *Variovorax* isolates in our SynCom. A minimal set of 7 of these genes (*iacABCDEFI*) was shown to be necessary and sufficient for auxin degradation^18^. We identified homologs for these genes across the *Variovorax* phylogeny (Extended Figure 5a) at relatively low sequence identity (27-48%). Two genes of the minimal set of 7 genes did not have any homologs in most *Variovorax* genomes (*iacB* and *iacI*). In addition to the *iac* operon, we scanned the genomes for the auxin degradation operon described by Ebenau-Jehle et al^63^ and could not identify it in any of the *Variovorax* isolates.

We also searched for the ACC deaminase gene by looking for the KEGG orthology id K01505 (1-aminocyclopropane-1-carboxylate deaminase) across the IMG annotations available for all our genomes.

### 19. *Variovorax* CL14 RNA-Seq in monoculture and in co-culture with *Arthrobacter* CL28

#### a. Bacterial culture

*Variovorax* CL14 was grown either alone or in co-culture with *Arthrobacter* CL28 in 5mL of 1/10 2xYT medium (1.6 g/L tryptone, 1 g/L yeast extract, 0.5 g/L NaCl) in triplicate. The mono-culture was inoculated at OD_600_ of 0.02 and the co-culture was inoculated with OD_600_ of 0.01 of each strain. Cultures were grown at 28°C to early stationary phase (approximately 22 hours) and cells were harvested by centrifugation at 4100 x g for 15 min and frozen at -80°C prior to RNA extraction.

#### b. RNA extraction and RNA-Seq

Cells were lysed for RNA extraction using TRIzol Reagent (Invitrogen) according to the manufacturer instructions. Following cell lysis and phase separation, RNA was purified using the RNeasy Mini kit (Qiagen) including the optional on column DNase Digestion with the RNAse-Free DNase Set (Qiagen). Total RNA quality was confirmed on the 2100 Bioanalyzer instrument (Agilent) and quantified using a Qubit 2.0 fluorometer (Invitrogen). RNA-Seq libraries were prepared using the Universal Prokaryotic RNA-Seq, Prokaryotic AnyDeplete kit (Tecan, formerly NuGEN). Libraries were pooled and sequenced on the Illumina HiSeq4000 platform to generate 50-bp single-end reads.

#### c. RNA-Seq analysis

We mapped the generated raw reads to the *Variovorax* CL14 genome (fasta file available on github.com/isaisg/variovoraxRGI) using bowtie2 with the ‘very-sensitive’ flag. We then counted hits to each individual coding sequence (CDS) annotated for the *Variovorax* CL14 genome using the function featureCounts from the R package Rsubread, inputting the *Variovorax* CL14 gff file (available on github.com/isaisg/variovoraxRGI) and using the default parameters with the flag allowMultiOverlap=FALSE. Finally, we used DESeq2 to estimate differentially expressed genes (DEGs) between treatments with the corresponding fold change estimates and false discovery adjusted p-values.

### 20. *Variovorax* CL14 genomic library construction and screening

#### a. Library construction

High molecular weight *Variovorax* CL14 genomic DNA was isolated by phenolchloroform extraction. This genomic DNA was partially digested with Sau3A1 (New England Biolabs), and separated on the BluePippin (Sage Science) to isolate DNA fragments >12.5 kb. Vector backbone was prepared by amplifying pBBR-1MCS2^64^ using Phusion polymerase (New England Biolabs) with primers JMC277-JMC278 (Supplementary Table 19), digesting the PCR product with BamHI-HF (New England Biolabs), dephosphorylating with Quick CIP (New England Biolabs), and gel extracting using the QIAquick Gel Extraction Kit (Qiagen). The prepared *Variovorax* CL14 genomic DNA fragments were ligated to the prepared pBBR1-MCS2 vector backbone using ElectroLigase (New England Biolabs) and transformed by electroporation into NEB 10-beta Electrocompetent *E. coli* (New England Biolabs). Clones were selected by blue-white screening on LB plates containing 1.5% agar, 50 µg/mL kanamycin, 40 µg/mL X-gal (5-bromo-4-chloro-3-indolyl-β-D-galactopyranoside), and 1 mM Isopropyl β-d-1-thiogalactopyranoside (IPTG) at 37°C. White colonies were screened by colony PCR using Taq polymerase and JMC247-JMC270 primers (Supplementary Table 19) to eliminate clones with small inserts. The screened library clones were picked into LB media + 50 µg/mL kanamycin, grown at 37°C, and stored at -80°C in 20% glycerol. The *Variovorax* CL14 genomic library comprises approximately 3,500 clones with inserts >12.5 kb in vector pBBR1-MCS2 in NEB 10-beta *E. coli*.

#### b. Library screening for IAA degradation

To screen the *Variovorax* CL14 genomic library for IAA degradation, the *E. coli* clones were grown in LB media containing 50 µg/mL kanamycin, 1 mM IPTG, 0.05 mg/mL IAA, and 0.25% ethanol from IAA solubilization for 3 days at 37°C. Salkowski reagent (10 mM FeCl_3_ in 35% perchloric acid) was mixed with culture supernatant 2:1 and color was allowed to develop for 30 min prior to measuring the absorbance at 530nm. Two clones from the library (plate 8 well E8 and plate2 well F10, henceforth Vector 1 and Vector 2, respectively) were identified as degrading IAA. The *Variovorax* CL14 genes contained in Vectors 1 and 2 were inferred by isolating plasmid from these clones using the ZymoPURE II Plasmid Midiprep Kit (Zymo Research) and Sanger sequencing the insert ends using primers JMC247 and JMC270 (Supplementary Table 19). Double digest of the purified plasmids with SacI and EcoRV confirmed the size of the inserts. Vector 1 contains a 35 kb insert and Vector 2 contains a 15kb insert (nucleotide coordinates 29100-64406 and 52627-67679, respectively, from *Variovorax* CL14 scaffold Ga0102008_10005 (Fig. 5a). The genes in both inserts are in the same direction as the IPTG inducible *Lac* promoter used to drive *LacZ*-alpha expression for blue-white screening on pBBR1-MCS2.

21. *Acidovorax* Root219::EV and *Acidovorax* Root219::V2 construction and screening Triparental mating was used to mobilize Vector 2 or control empty vector (EV) pBBR1-MCS2 from *E. coli* to *Acidovorax* Root219. Donor NEB 10-beta *E. coli* containing the vector for conjugation and helper strain *E. coli* pRK2013^65^ were grown in LB medium containing 50 µg/mL kanamycin at 37°C. *Acidovorax* Root219 was grown in 2xYT medium containing 100 µg/mL ampicillin at 28°C. Bacteria were washed 3 times with 2xYT medium without antibiotics, mixed in a ratio of approximately 1:1:10 donor:helper:recipient, centrifuged and resuspended in 1/10 the volume and plated as a pool on LB agar plates without antibiotics and grown at 28°C. 18-30 hours later, exconjugantes were streaked on LB agar plates containing 50 µg/mL kanamycin and 100 µg/mL ampicillin to select only *Acidovorax* Root219 containing the conjugated vector. The resulting strains are designated *Acidovorax* Root219-EV containing empty vector pBBR1-MCS2 and *Acidovorax* Root219-V2 containing Vector 2 (Methods 20b). *In vitro* IAA degradation was performed as in Methods 14 using M9 media with carbon sources: 15 mM succinate, 0.1 mg/mL indole-3-acetic acid (IAA), and 0.5% Ethanol with the addition of 50 µg/mL kanamycin and 1mM IPTG. Primary root elongation measurement was performed as in Methods 2c on MS medium with 1mM IPTG and RGI induced by either 1µM IAA or *Arthrobacter* CL28.

### 21. *Variovorax* hotspot 33 knockout construction and screening

The unmarked deletion mutant *Variovorax* CL14 Δ2643613653-2643613677 (*Variovorax* CL14ΔHS33) was constructed based on a genetic system developed for *Burkholderia* spp. and its suicide vector pMo130^66^

#### a. Knockout suicide vector pJMC158 construction

The vector backbone was amplified from pMo130 using primers JMC203-JMC204 (Supplementary Table 19) with Q5 DNA polymerase (New England Biolabs), cleaned up and treated with DpnI (New England Biolabs). 1Kb regions for homologous recombination flanking *Variovorax* CL14 genes 2643613653-2643613677 were amplified using Q5 Polymerase (New England Biolabs) and primers JMC533-JMC534 and JMC535-JMC536 (Supplementary Table 19). The vector was assembled with Gibson Assembly Mastermix (New England Biolabs) at 50°C for 1 hour, transformed into NEB 5-alpha chemically competent *E. coli* (New England Biolabs), and plated on LB agar with 50 µg/mL kanamycin. pJMC158 DNA was isolated from a clone using the ZR Plasmid Miniprep Classic Kit (Zymo Research), sequence confirmed, and transformed into biparental mating strain *E. coli* WM3064. *E. coli* strain WM3064 containing pJMC158 was maintained on LB containing 50 µg/mL kanamycin and 0.3mM diaminopimelic acid (DAP) at 37°C.

#### b. Conjugative transfer of pJMC158 into *Variovorax* CL14

For biparental mating, *E. coli* WM3064 was grown as above, and *Variovorax* CL14 was grown in 2xYT medium containing 100 µg/mL ampicillin. Each strain was washed separately 3 times with 2xYT medium, then mixed at ratios between 1:1-1:10 donor:recipient, centrifuged and resuspended in approximately 1/10 the volume and plated in a single pool on LB agar containing 0.3mM DAP and grown at 28°C overnight. Exconjugants were streaked onto LB plates containing 100 µg/mL ampicillin 50 µg/mL kanamycin lacking DAP and grown at 28°C to select *Variovorax* CL14 strains that incorporated suicide vector pJMC158. First crossover strains were subsequently purified once by re-streaking and then individual colonies grown in LB with 100 µg/mL ampicillin 50 µg/mL kanamycin.

#### c. Resolution of pJMC158 integration and knockout strain purification and verification

To resolve the integration of pJMC158, first crossover strains were grown once in LB medium containing 100 µg/mL ampicillin and 1 mM IPTG then plated on media containing 10 g/L tryptone, 5 g/L yeast extract, 100 g/L sucrose, 1.5% agar, 100 µg/mL ampicillin and 1 mM IPTG. Colonies were picked into the same liquid media and grown once. The resulting strains were screened by PCR using Q5 polymerase for deletion of genes 2643613653-2643613677 using primers JMC568-JMC569 (Supplementary Table 19). These strains were subsequently plate purified at least 3 times on LB 100 µg/mL ampicillin plates. To ensure strain purity, PCR primers were designed to amplify from outside into the genes that were deleted (primer pairs JMC571-JMC569 and JMC568-JMC570, Supplementary Table 19). These PCR reactions were performed using Q5 polymerase with wild type *Variovorax* CL14 as a control. All genomic DNA used for screening PCR was isolated using the Quick-DNA miniprep kit (Zymo Research). The resulting knockout strain was designated *Variovorax* CL14ΔHS33.

#### d. Screening of *Variovorax* CL14 ΔHS33

*In vitro* IAA degradation was performed as in Methods 14 using M9 media with carbon sources: 15 mM succinate, 0.1 mg/mL indole-3-acetic acid (IAA), and 0.5% Ethanol. Primary root elongation measurement was performed as in Methods 2c on MS medium with 1mM IPTG and RGI induced by either 1µM IAA or *Arthrobacter* CL28.

## Acknowledgments

We thank Stratton Barth, Julia Shen, May Priegel, Dilan Chudasma, Darshana Panda, Izabella Castillo, Nicole Del Risco, and Chloe Lindberg for technical assistance; Dale Pelletier, DOE-ORNL and Paul Schulze-Lefert, MPIPZ, Koeln, Germany for strains; the Dangl lab microbiome group for useful discussions; Anna Stepanova, Jose Alonso, Javier Brumos (North Carolina State University, USA), Joseph Kieber, Jason Reed (UNC Chapel Hill), and Isaac Greenhut (University of California, Davis) for useful discussions and Derek Lundberg (Max Planck Institute for Developmental Biology, Tübingen, Germany), and Anthony Bishopp (University of Nottingham, UK) for critical comments on the manuscript.

## Author contributions

Conceptualization: O.M.F., I.S.G, G.C., J.L.D.; Methodology: O.M.F., I.S.G, G.C., T.F.L., J.M.C., P.J.P.L.T., E.D.W., C.R.F.; Software: I.S.G.; Validation: O.M.F., I.S.G, G.C., T.F.L., J.M.C.; Formal analysis: I.S.G, O.M.F., G.C.; Investigation: O.M.F., I.S.G, G.C., T.F.L., J.M.C., P.J.P.L.T., E.D.W., C.R.F; Resources: J.L.D.; Data Curation: I.S.G, O.M.F.; Writing – original draft: O.M.F, I.S.G, G.C., J.M.C.; Writing – review and editing: T.F.L.,, C.D.J., J.L.D.; Visualization: I.S.G, O.M.F., G.C.; Supervision: J.L.D, C.D.J.; Project administration: J.L.D.; Funding acquisition: J.L.D.

## Data and materials availability

The 16S rRNA amplicon sequencing data associated with this study was deposited in the NCBI SRA archive under the project accession PRJNA543313. The raw transcriptomic data was deposited in the Gene Expression Omnibus (GEO) under the accession GSE131158. In addition to the supplementary tables, we deposited all scripts and additional data structures required to reproduce the results of this study in the following GitHub repository: https://github.com/isaisg/variovoraxRGI

---

**Extended data Fig. 1.**
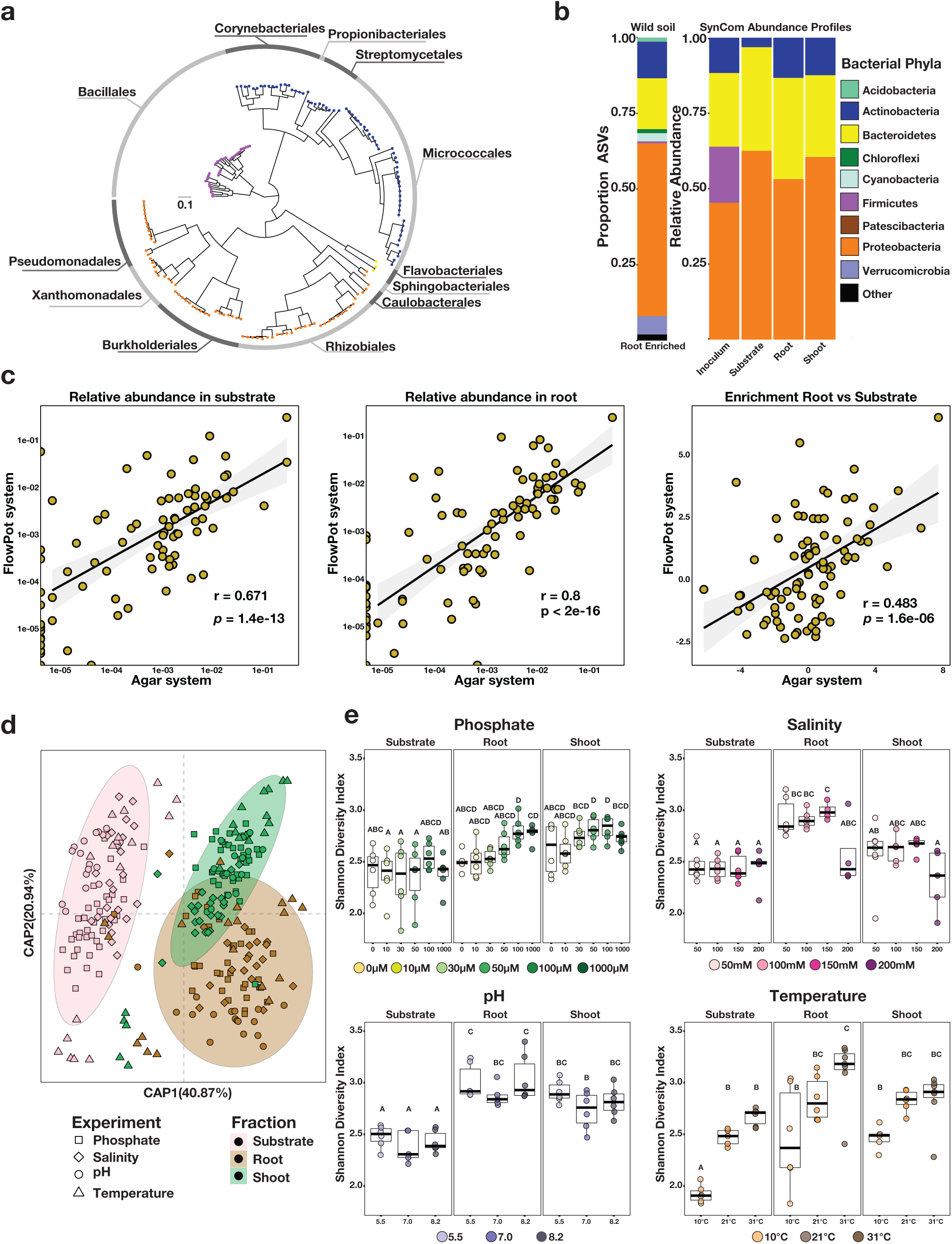
Synthetic community resembles the taxonomic makeup of natural communities. **(a)** Phylogenetic tree of 185 bacterial genomes included in the synthetic community (SynCom). The tree tips are colored according to phylum. The outer ring shows the distribution of the 12 distinct bacterial orders present in the SynCom. **(b)** The panel on the left (wild soil) shows the proportion of amplicon sequence variants (ASVs) enriched (q-value < 0.1) in the plant root in comparison to soil in a microbiota profiling study from the same soil that SynCom strains were isolated in Levy et al20. In the panel, ASVs are colored according to phylum and the ASVs belonging to Proteobacteria are colored by class. The panel on the right (SynCom panel) represents the relative abundance profiles of bacterial isolates across the initial inoculum, planted agar, root and shoot in plant exposed to the full SynCom. Bacterial isolates are colored based on their phylum, the Proteobacteria are colored according to the class level classification. **(c)** Comparison of SynCom community composition in the agar and soil-based microcosms. (Left) Relative abundance in the substrate (FlowPot system) and (middle) root, (right) as well as root vs substrate enrichment levels are shown. Each dot represents a single USeq. Pearson correlation line, 95% confidence intervals, r value and p value are shown for each comparison. **(d)** Canonical analysis of principal coordinates showing the influence of the fraction (planted agar, root, shoot) on the assembly of the bacterial SynCom across the four gradients used in this work (phosphate, salinity, pH, temperature). Different colors differentiate between the fractions and different shapes differentiate between experiments. Ellipses denote the 95% confidence interval of each fraction. **(e)** Abiotic conditions displayed reproducible effects on alpha-diversity. Each panel represents the bacterial alpha-diversity across the different gradient conditions (phosphate, salinity, pH, temperature) and the fractions (planted agar, root, shoot) used in this work. Bacterial alpha-diversity was estimated using Shannon Diversity. Letters represent the results of the post hoc test of an ANOVA model testing the interaction between fraction and abiotic condition.

**Extended data Fig. 2.**
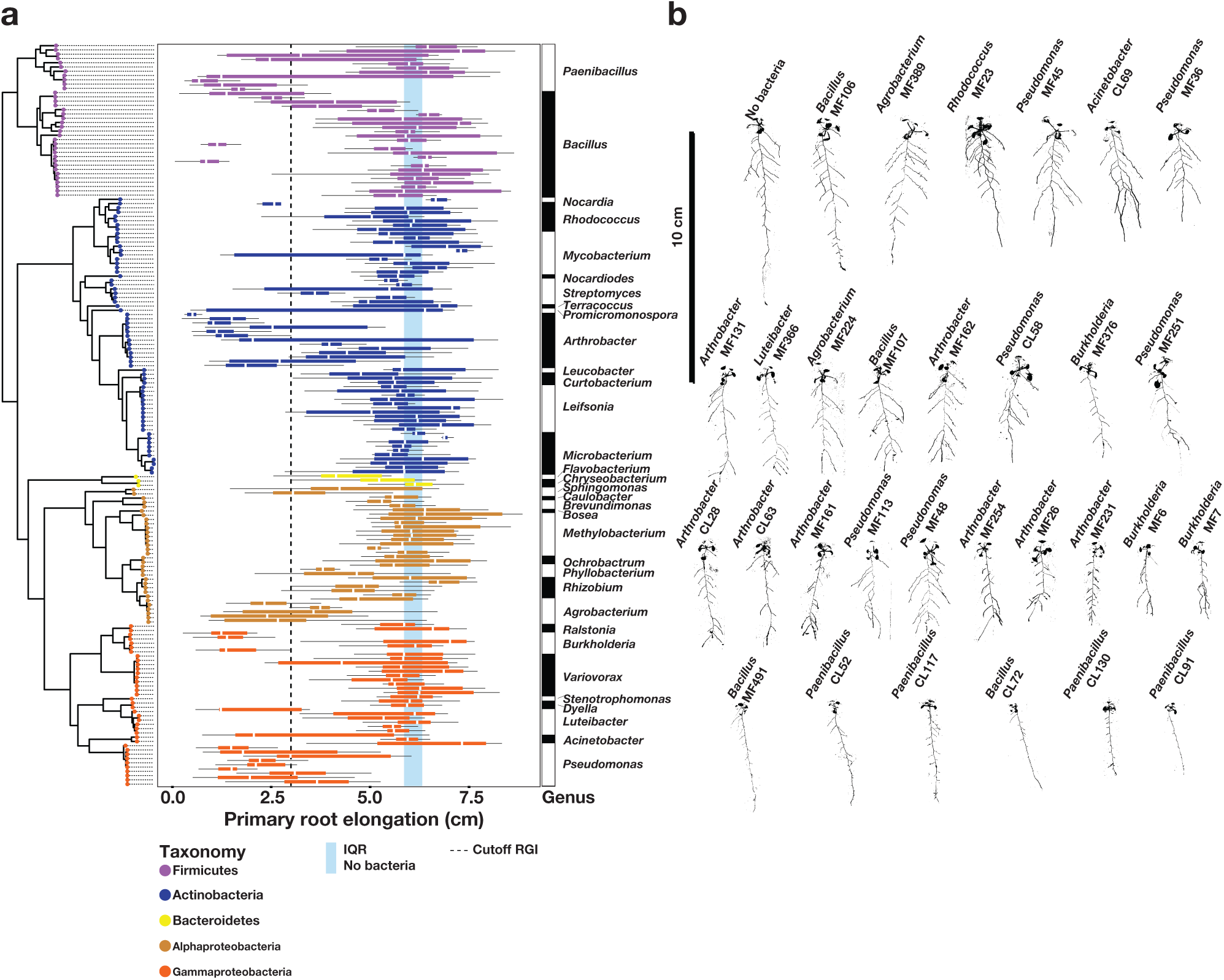
Root growth inhibition trait is distributed across bacterial phylogeny. **(a)** Primary root elongation of plants inoculated with single bacterial isolates (one boxplot per isolate). Isolates are ordered according to the phylogenetic tree on the left side of the panel and colored based on their genome-based taxonomy. The vertical blue strips across the panel corresponds to the interquartile range (IRQ) of plants grown in sterile conditions. The vertical dotted line represents the 3 cm cutoff used to classify strains as root growth inhibiting (RGI) strains. The bar on the right side of the panel denotes the genus classification of each isolate. **(b)** Binarized image of representative seedlings grown axenically (No bacteria) or with thirty-four RGI strains individually.

**Extended data Fig. 3.**
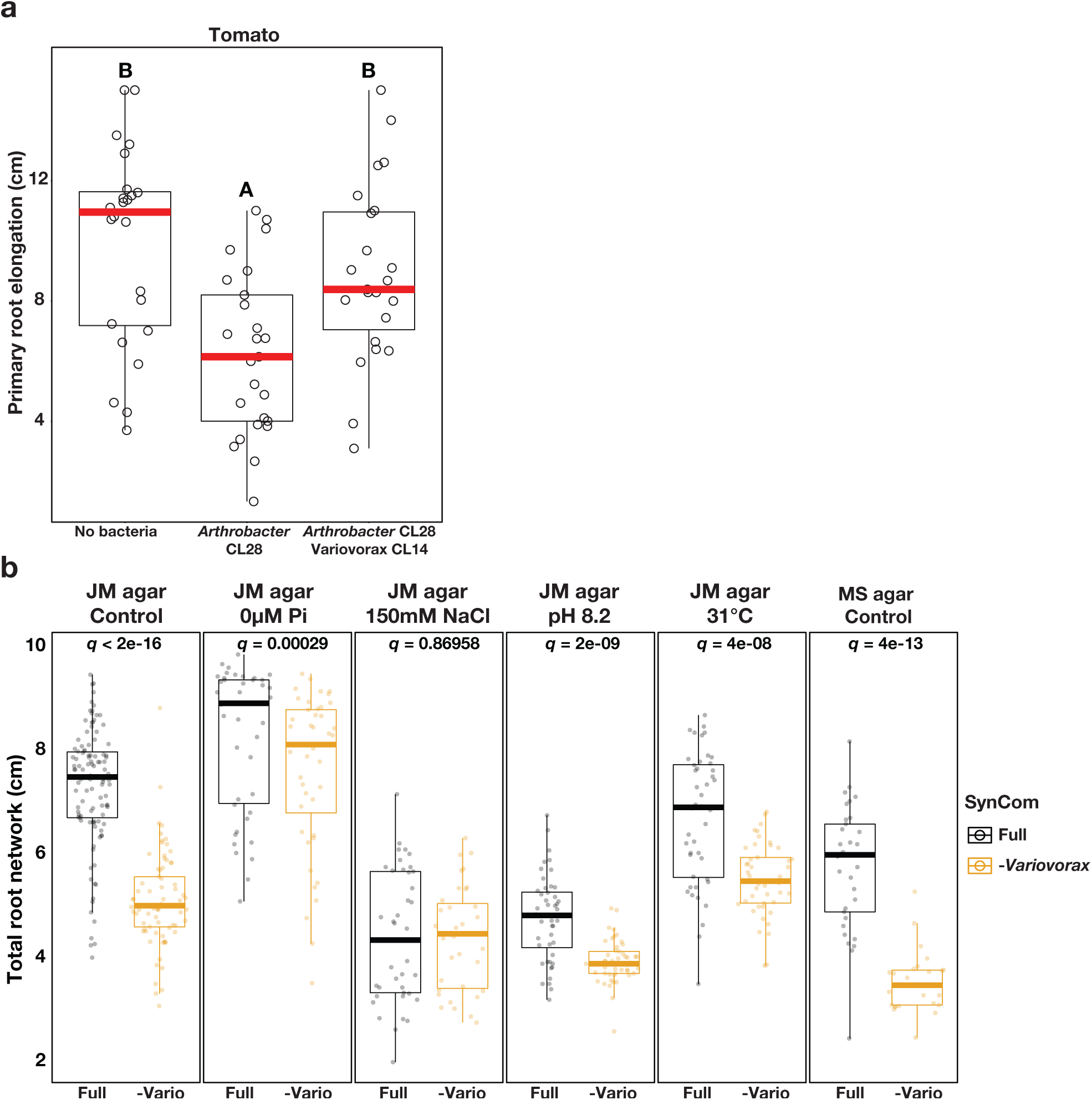
Variovorax-mediated reversion of root growth inhibition. **(a)** Variovorax-mediated reversion of root growth inhibition is maintained in a second plant species. Primary root elongation of uninoculated tomato seedlings (No bacteria) or seedlings inoculated with the Arthrobacter CL28 individually or along with Variovorax CL14. Letters indicate post hoc significance. **(b)** Total root network of Arabidopsis seedlings grown with the full SynCom (Full) or with the full SynCom excluding Variovorax (-Variovorax) across different abiotic conditions: full medium (JM agar control), phosphate starvation (JM agar 0 uM Pi), salt stress (JM agar 150 mM NaCl), high pH (JM agar pH 8.2) and high temperature (JM agar 31oC) and half Murashige and Skoog medium (MS agar control). Letters indicate statistical significance using ANOVA performed within each experimental condition.

**Extended data Fig. 4.**
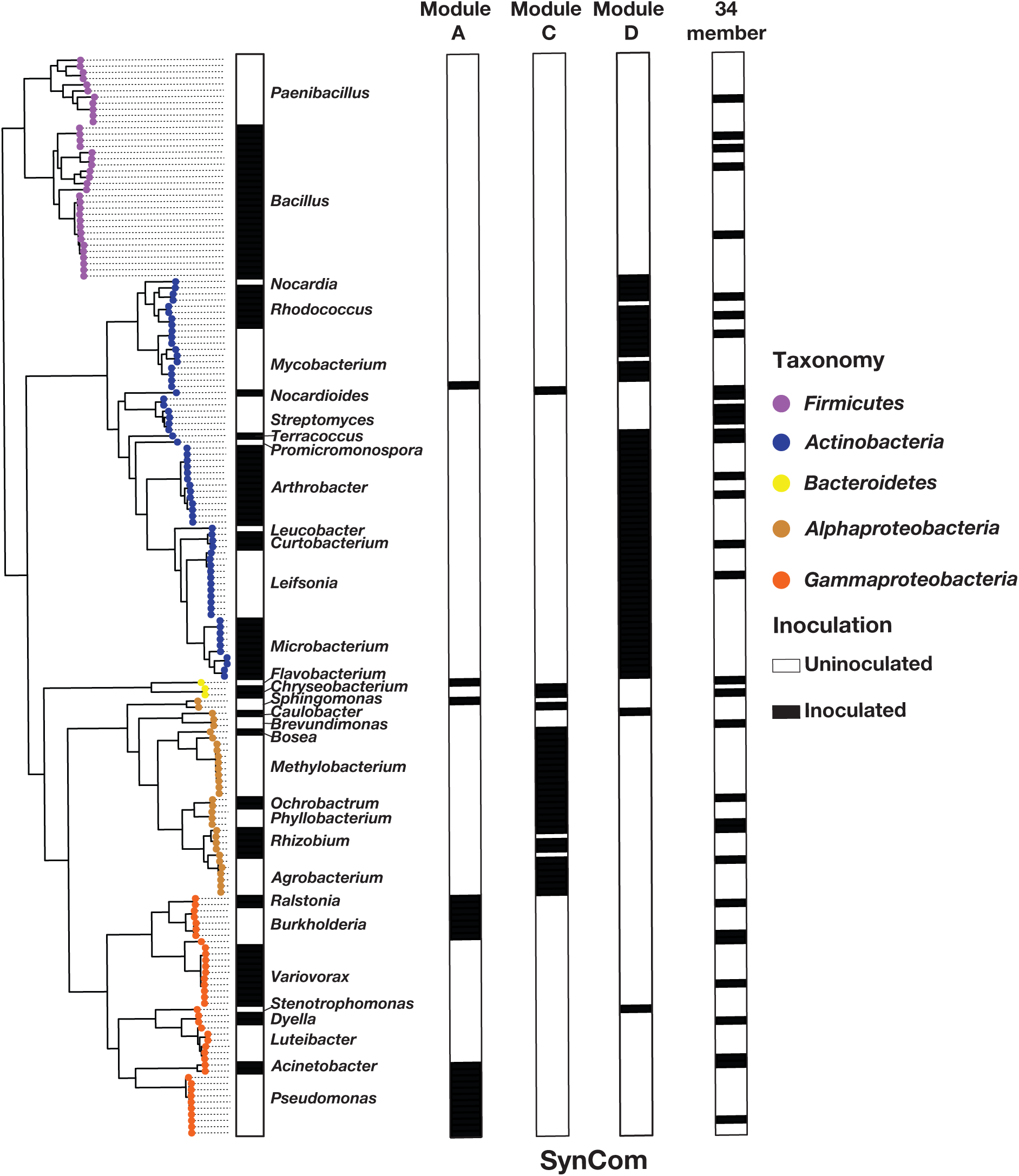
Taxonomic composition of the SynCom used in Figure 2D. Bar graphs showing the isolate composition of SynComs composed by module A (Module A), module C (Module C), module D (Module D) and a 34-member synthetic community (34 member)2. Isolates are ordered according to the phylogenetic tree on the left side of the panel. The tips of the phylogenetic tree are colored based on the genome-based taxonomy of each isolate. Presence of an isolate across the different SynComs is denoted by a black filled rectangle.

**Extended data Fig. 5.**
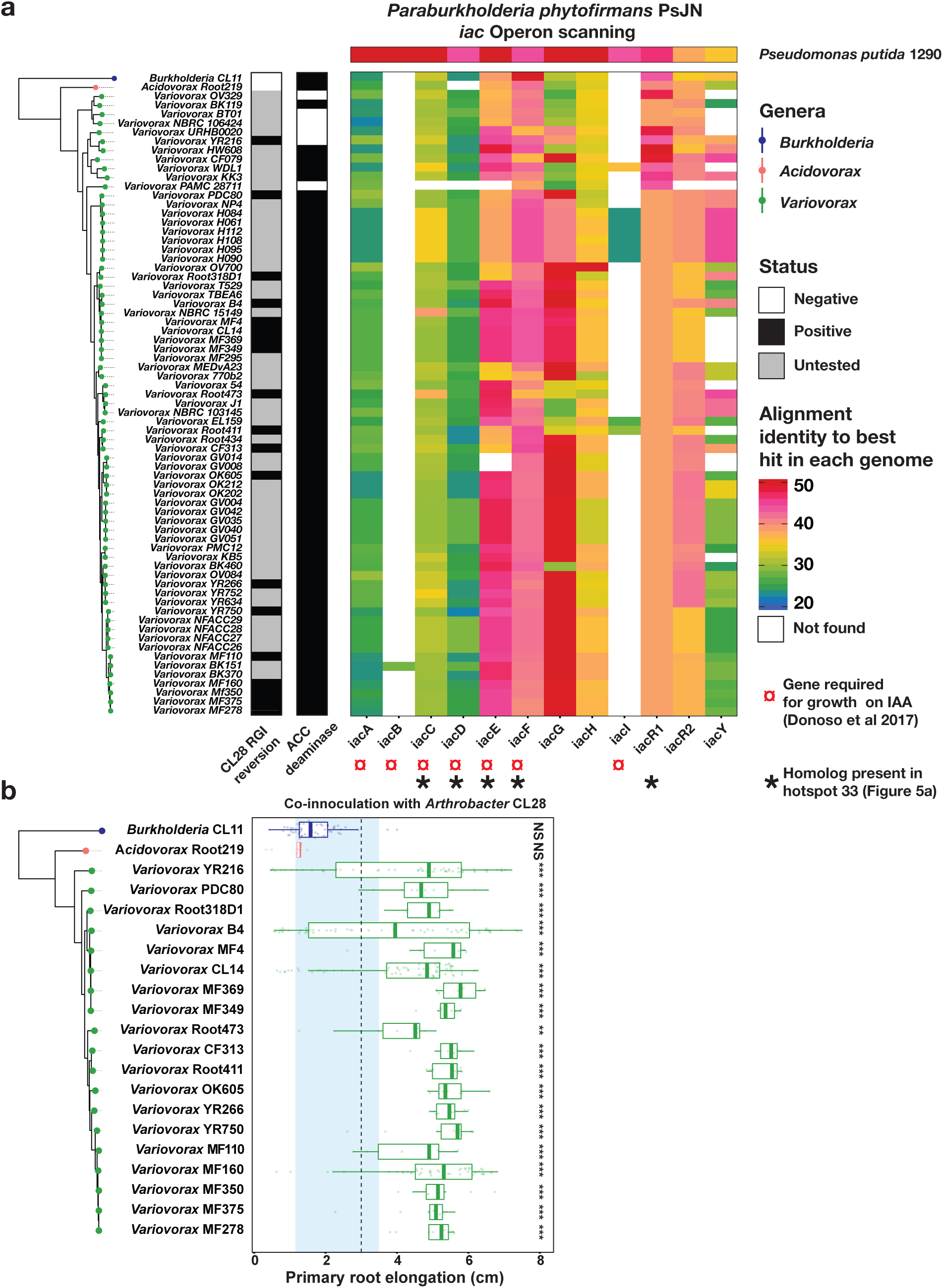
Reversion of root growth inhibition is prevalent across the Variovorax phylogeny. **(a)** Phylogenetic tree of 54 publically available Variovorax genomes and two outgroup isolates, Acidovorax Root219 and Burkholderia CL11. The CL28 RGI reversion bar binarizes (positive, negative, untested) the ability of each isolate in the phylogeny to revert the root growth inhibition caused by Arthrobacter CL28. The ACC deaminase bar denotes the presence of the KEGG orthology term KO1505 (1-aminocyclopro-pane-1-carboxylate deaminase) in each of the genomes. The heatmap denotes the percent identity of BLASTp hits in the genomes to the genes from the auxin degrading iac operon in Paraburkholderia phytophirmans described by Donoso, et al.33. Note that synteny is not conserved and these BLAST hits are spread throughout the genomes. **(b)** Phylogenetic tree of 19 Variovorax genomes along two outgroup isolates, Acidovorax Root219 and Burkholderia CL11 that were tested for their ability to revert the root growth inhibition (RGI) imposed by Arthrobacter CL28. The blue vertical strip across the panel denotes the interquartile range of plants treated solely with Arthrobacter CL28. The dotted vertical line across the panel denotes the 3 cm cutoff used to classify a treatment as a root growth inhibitor (RGI). Each boxplot is colored according to the genus classification of each isolate. Statistical significance is denoted on the top of each boxplot.

**Extended data Fig. 6.**
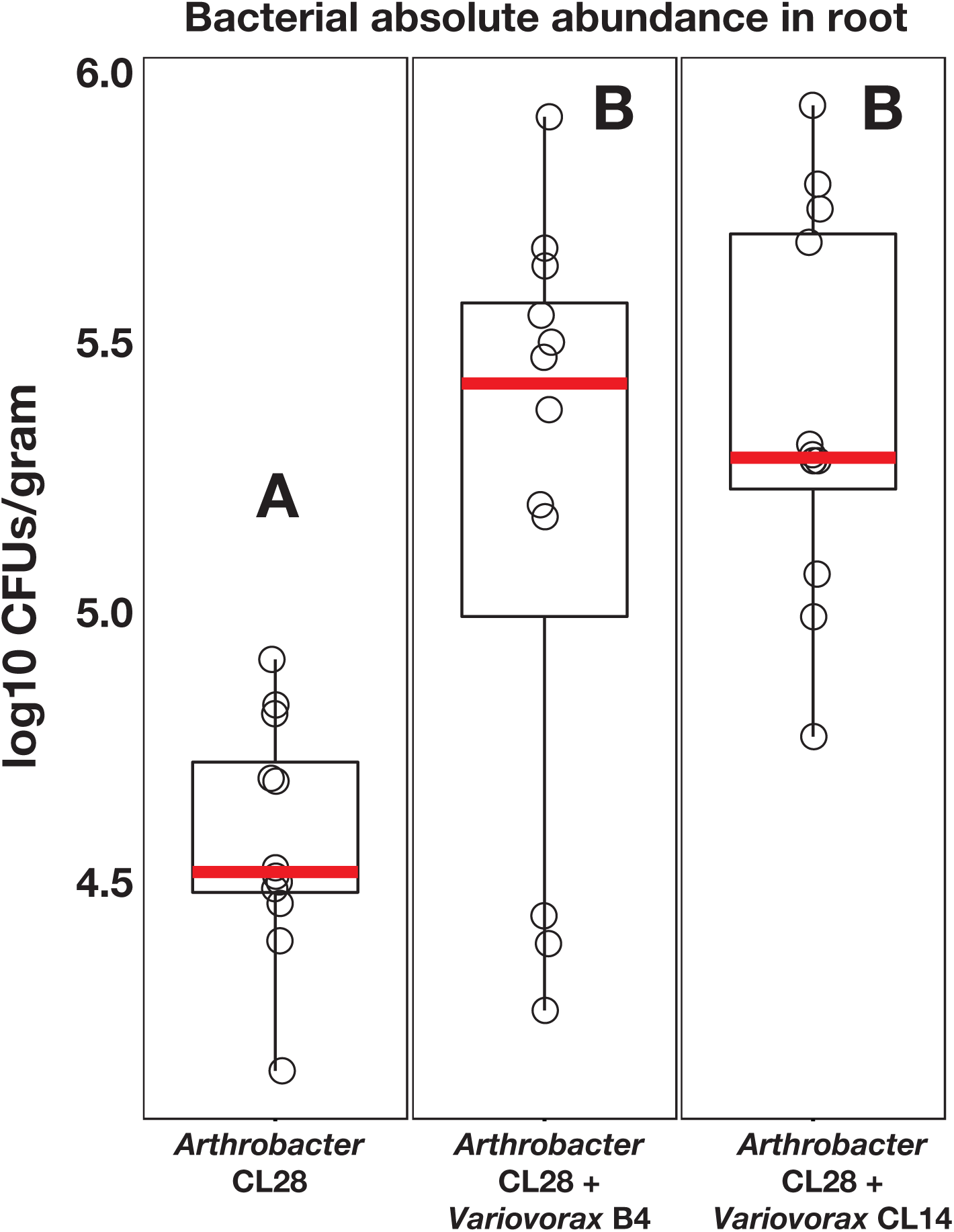
Variovorax does not inhibit the growth of an RGI strain. In planta absolute abundance of Arthrobacter CL28 when inoculated alone or with two Variovorax representatives: Variovorax B4 and Variovorax CL14. Log-transformed-Colony forming Units (CFU) of Arthrobacter CL28 normalized to root weight are shown. To selectively grow Arthro-bacter CL28, CFUs were counted on Luria Bertani (LB) agar plates containing 50 µg/ml of Apramycin, on which both Variovorax B4 and Variovorax CL14 do not grow.

**Extended data Fig. 7.**
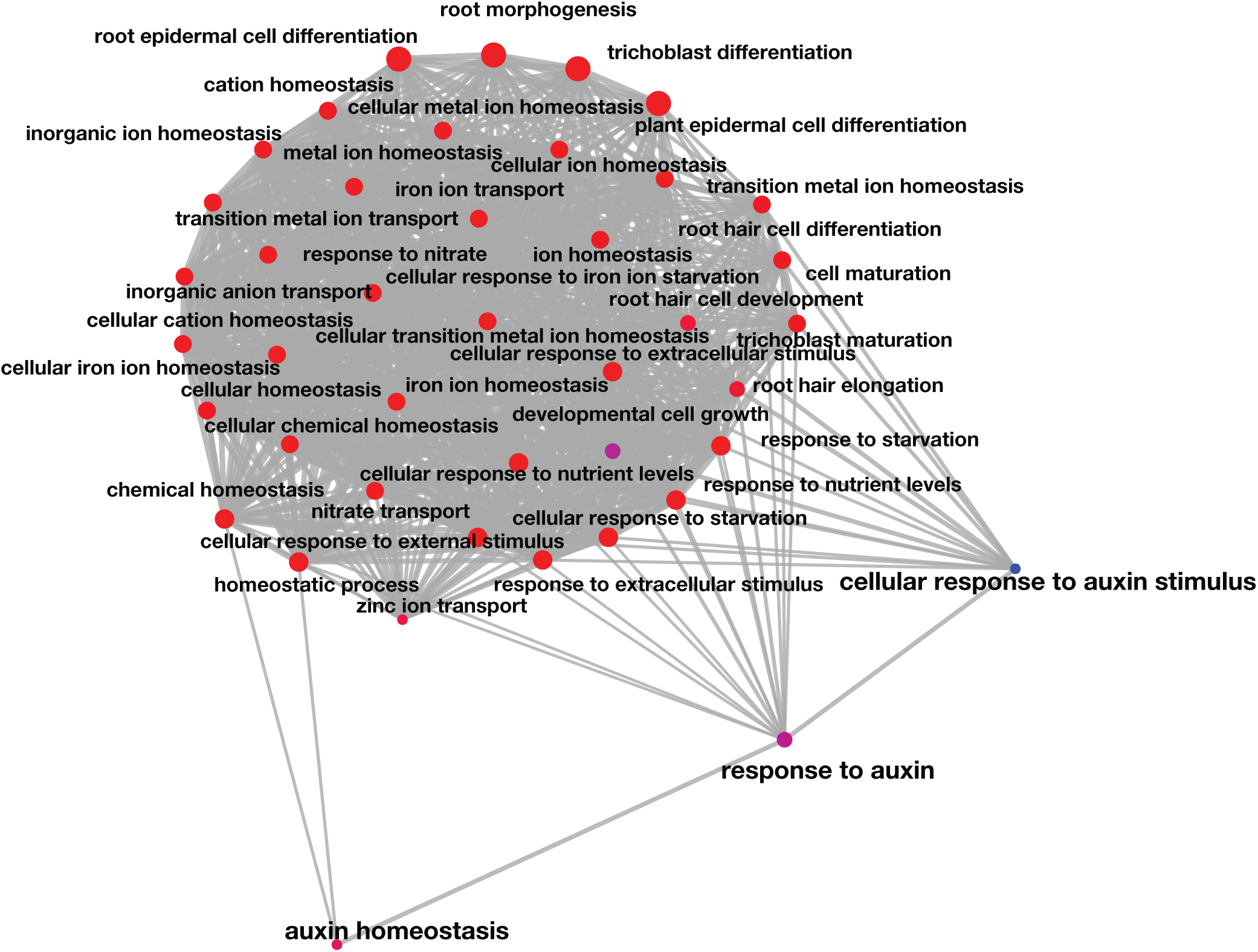
Root growth inhibition-related genes share gene ontologies. Network of statistically significant gene ontology terms contained in the 18 genes upregulated in Variovorax CL14/Arthrobacter CL28 co-inoculation vs Arthrobacter CL28 alone AND in the full SynCom vs the Variovorax drop-out SynCom (See Figure 4a and 4b). The network was computed using the emapplot function from the package clusterProfiler in R. A p-value for terms across the gene ontology was computed using a hypergeometric test, additionally the size of each point (Gene ontology term) denoted the number of genes mapped in that particular term.

**Extended data Fig. 8.**
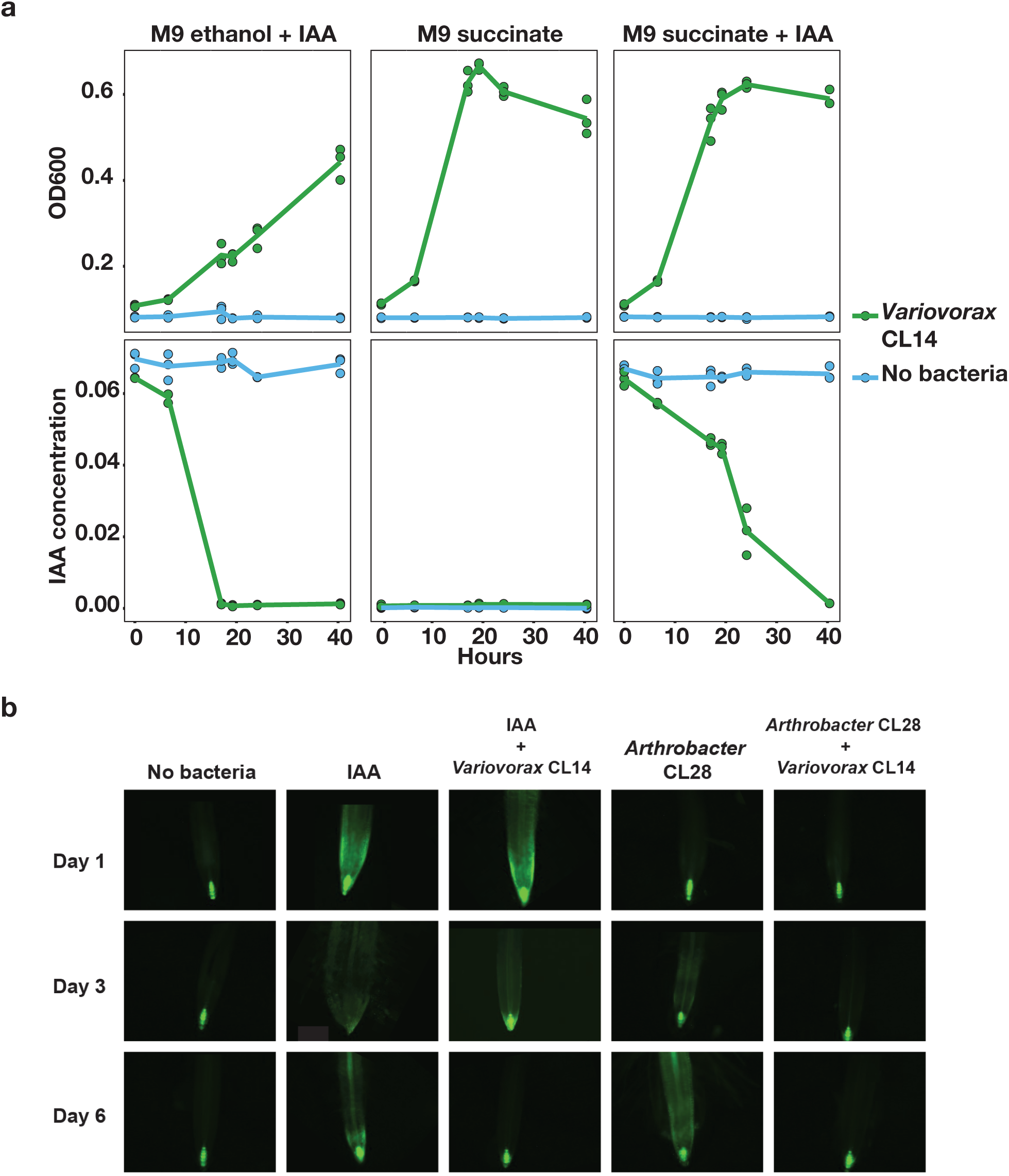
Variovorax degrades auxin and quenches auxin perception by the plant. **(a)** Variovorax utilizes auxin as a carbon source. Growth curves showing Optical density at OD600 (top) and Indole-3-acetic acid (IAA) concentrations (mg/mL bottom) in Variovorax CL14 cultures grown in M9 media with carbon sources: IAA and ethanol (from IAA solubilization) (left), succinate (center), and succinate, IAA, and ethanol (right). **(b)** Variovorax quenches auxin bioreporter DR5::GFP induction. Main root tips of DR5::GFP plants grown with different Indole-3-acetic acid (IAA) and bacterial treatments. DR5::G-FP plants were treated with the tripartite system (Arthrobacter CL28, Variovorax CL14, CL28+CL14), with the drop-out System (full Syncom, Variovorax drop-out Syncom [-Variovorax]), IAA and IAA+Variovorax CL14. GFP fluorescence was imaged 1, 3 and 6 days post inoculation. Fluorescence was quantified in the root elongation zone.

**Extended Data Fig. 9.**
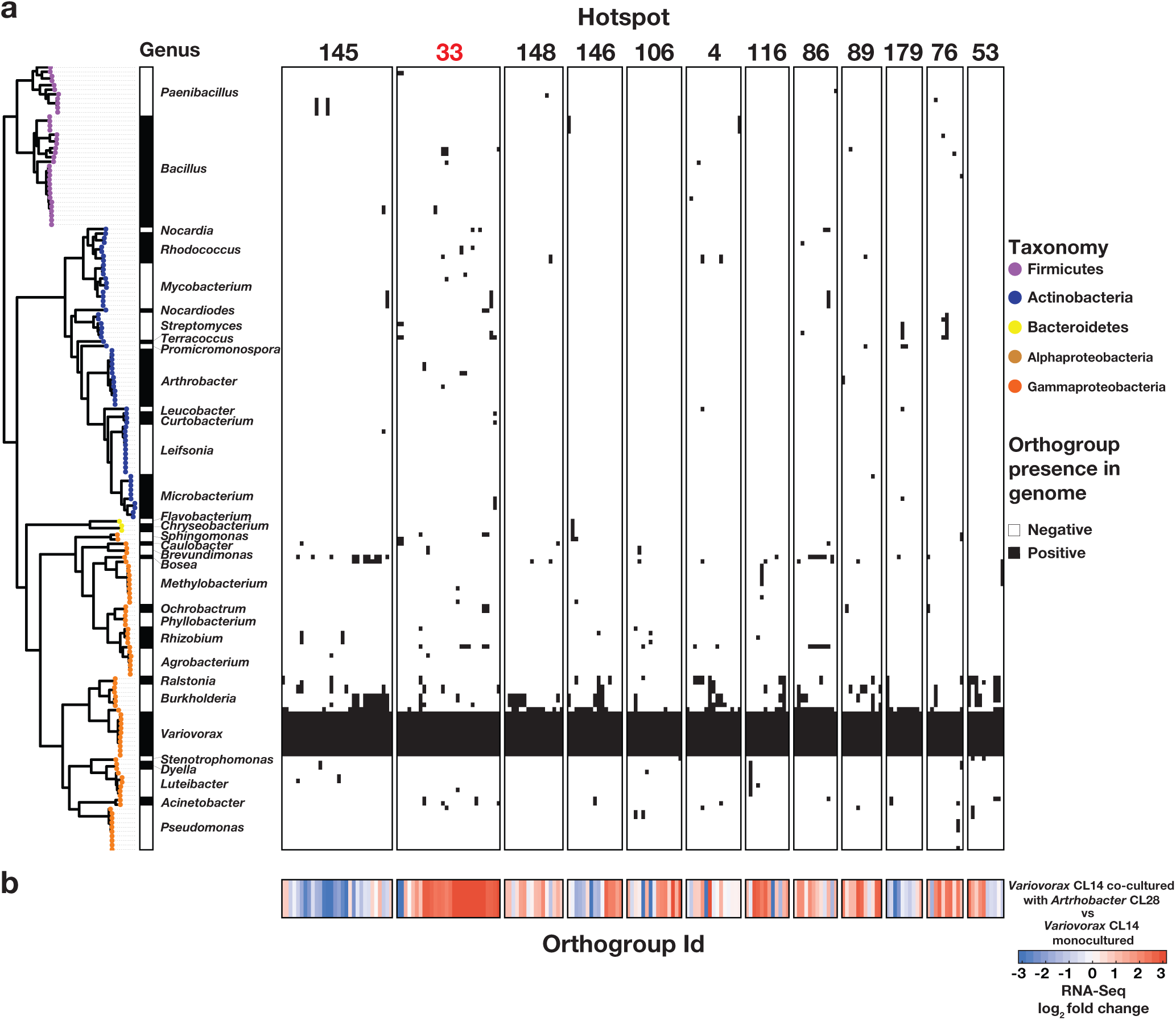
Detection of CL28-responsive Variovorax-unique operons. **(a)** Presence/absence matrix denoting the distribution of 12 Variovorax-unique hotspots containing at least 10 genes across the 185 members of the SynCom. Hotspots are defined using the Variovorax CL14 genome as a reference. Phylogeny of the 185 SynCom members is shown to the left of the matrix. We determined the presence of an orthogroup based on a hidden Markov model profile scanning of each orthogroup across the 185 genomes in the SynCom. **(b)** Results of bacterial RNA-Seq. Log2(fold change) of each gene shown in the matrix in Variovorax CL14 is co-cultured with Arthrobacter CL28 versus Variovorax CL14 monoculture. Note uniform up-regulation of genes in cluster 33.

**Extended data Fig. 10.**
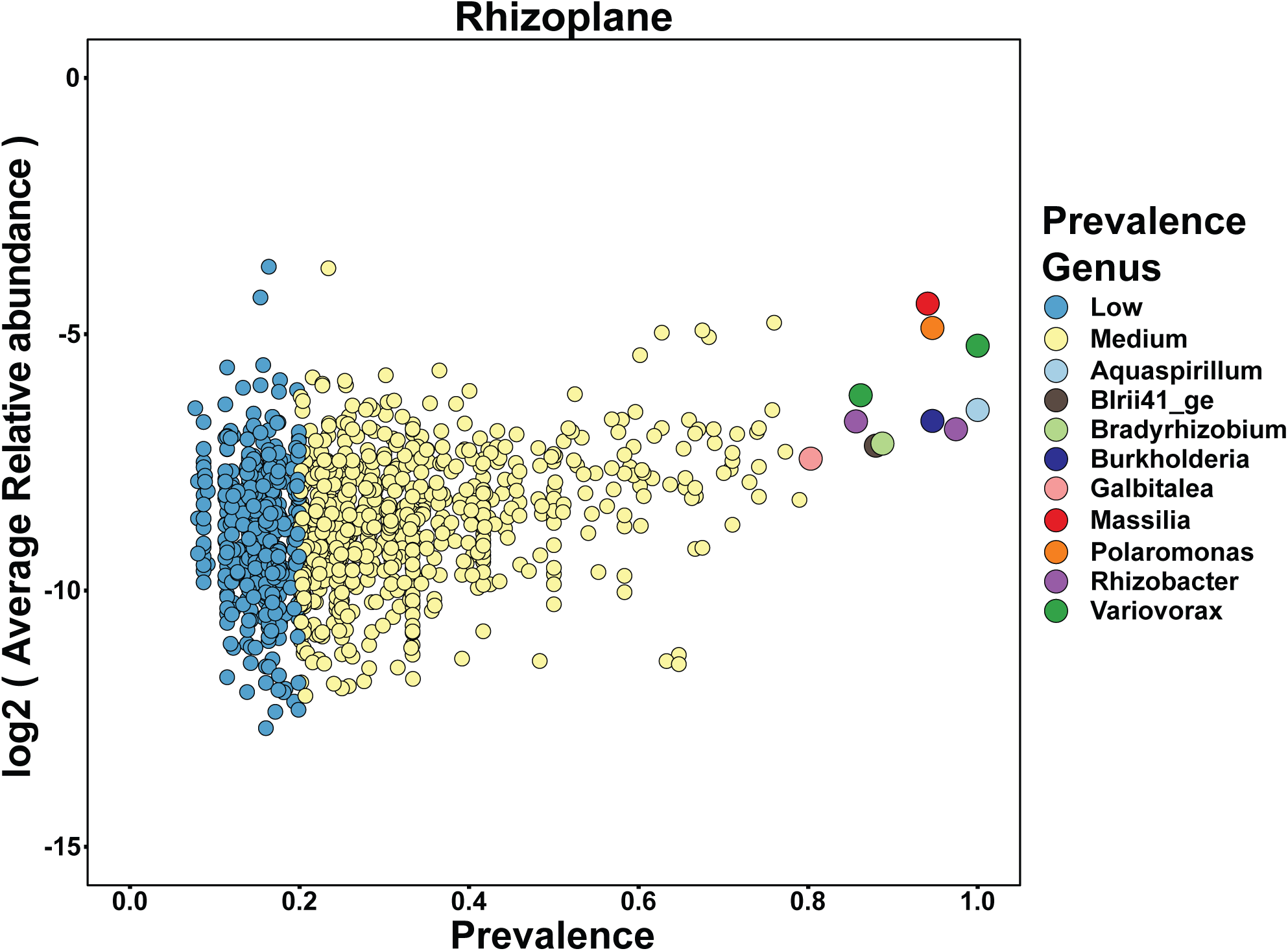
Variovorax are highly prevalent across naturally occurring Arabidopsis microbiomes. Correlation plot of data reanalyzed from Thiegart et al38 comparing bacterial amplicon sequence variants (ASV) prevalence to log transformed relative abundance in Arabidopsis thaliana rhizosphere samples taken across 3 years in 17 sites in Europe.

